# Functional impact of a type VI secretion system on RsmE-mediated spatial structure formation in *Pseudomonas fluorescens* Pf0-1

**DOI:** 10.64898/2026.07.29.741521

**Authors:** Meghan K. Wells, Anton F. Evans, Sadhana Srinivasa, Amber Delprince, Raziel Santos, Cole Garland, Anthony Valletta, Wook Kim

## Abstract

Bacteria naturally form densely structured populations wherein both space and nutrients become locally limited. In densely populated *Pseudomonas fluorescens* Pf0-1 colonies, spatiogenetic patches with drastically reduced cellular density naturally emerge through spontaneous mutations in *rsmE*. RsmE is a posttranscriptional regulator that binds to specific mRNAs, including those that consequently repress the production of extracellular secretions. We have previously shown that Δ*rsmE* produces an extracellular polysaccharide (EPS) and biosurfactant that collectively function to define the spatiogenetic structure and locally outcompete the WT. Here, we identify additional RsmE-regulated secretions through a combination of RNA-sequencing and LC-MS/MS. In particular, a type VI secretion system (T6SS) was exclusively produced by Δ*rsmE*. Confocal microscopy imaging of WT and genetically modified Δ*rsmE* cocultures showed that the T6SS kills WT cells that invade the low-density spatiogenetic structure. However, its impact on competition with the WT was both quantifiably and spatially limited in the absence of the normally co-produced EPS and biosurfactant, largely due to the consequently altered spatial structure. These results highlight the importance of understanding both the individual and collective spatial functions of extracellular secretions, especially in the ecological context of densely structured microbial populations.

**IMPORTANCE:** In crowded wild type (WT) *Pseudomonas fluorescens* Pf0-1 colonies, *rsmE* mutants spontaneously emerge by forming a low cellular density spatial structure that is devoid of WT cells. RsmE functions primarily to repress the production of various extracellular secretions, including an extracellular polysaccharide and biosurfactant, that collectively form the unique spatial structure. Here, we describe an RsmE-regulated type VI secretion system that kills the invading WT cells and protect the low-density structure. Importantly, individual RsmE-regulated secretions carry out unique functions, but their efficacies largely depend on other co-regulated products. Extracellular secretions that are mechanistically similar to those described here are co-produced across diverse biofilm-forming and virulent bacterial species, where they also likely play complex molecular and ecological roles.

## INTRODUCTION

Bacteria form biofilms on virtually any biotic or abiotic surfaces, wherein individual cells of the same and different genotypes interact chemically and physically in close proximity. A universal feature of biofilms is that the entire community is encased within an extracellular matrix (ECM), which forms dynamically through the localized accumulation of diverse compounds secreted by individual cells (1–5). Given the protective nature of the ECM against predation, dehydration, and lethal chemical infiltration, biofilms are often generalized to be a conserved differentiation program rooted by cooperating individuals (6–8). However, ECM production is also described as an adaptive response among the competing genotypes within a common space (5, 9–11). Although the ECM components of individual biofilms typically fall into four common categories – carbohydrates, proteins, nucleic acids, and lipids – the specific molecular composition could vary across different genotypes, species, and environmental conditions (2, 4, 5, 9). Many questions remain on the functional role of each matrix component, in particular, how they function together, if at all.

Experimental evolution studies of *Pseudomonas fluorescens* biofilms demonstrate that altered production of extracellular secretions lead to the emergence of striking social phenotypes. For example, mutations that elevate the production of aggregative extracellular secretions drive niche separation or cooperative colony expansion, while those that elevate the production of mucoid extracellular secretions drive spatial competition (12–18). In the latter case, diverse mutations in the *rsmE* gene produce a highly advantageous spatial phenotype that is specific to a crowded population (12). RsmE is an RNA-binding post-transcriptional regulator, and the observed mutations appear to de-repress the production of at least two visible extracellular secretions, a polysaccharide composed primarily of glucose (14) and a biosurfactant known to be the cyclic lipopeptide gacamide A (16, 19). Depending on the specific mutation in *rsmE*, the production of the biosurfactant and the degree of fitness vary relative to the parent *P. fluorescens* Pf0-1 strain (16). The two secretions function collectively to create and protect the spatiogenetic structure within a crowded population, where the polysaccharide functions to push away neighboring cells to create space and the biosurfactant appears to physically sequester the polysaccharide, and potentially additional secretions, to maintain a defined genotypic boundary (16). However, knocking out both secretions in a Δ*rsmE* mutant retained its ability to outcompete the parent strain, albeit at a significantly reduced level, indicating that additional secretions contribute to spatial structure formation (16).

*P. fluorescens* possesses two paralogs of RsmE – RsmA and RsmI – that do not influence spatial structure formation (16). In addition, the ability of the Δ*rsmE* mutant to outcompete the parent strain manifests exclusively when co-cultured in a structured population, indicating that RsmE-associated extracellular secretions specifically function to increase fitness through spatial interaction (12, 16). However, it remains unclear which RsmE-specific mRNA targets lead to these unique spatial interactions. Previous studies have identified hundreds of potential direct mRNA-binding partners of Rsm homologs in *Escherichia coli* (20–23), *Pseudomonas putida* (24), and *Pseudomonas aeruginosa* (23). These studies suggest that Rsm paralogs bind to a variety of mRNA leading to direct and indirect regulation of diverse cellular functions including metabolism, transport and secretion, stress response, and transcription regulation, among others (20–24). However, the number of direct mRNA binding partners of the Rsm paralogs varies greatly between different studies and organisms, as single omics studies are often limited by specific experimental conditions (21, 22). While these studies provide a starting point for understanding Rsm regulation, it remains unclear which of the hundreds of previously identified mRNA targets are both RsmE-specific and responsible for spatial interactions within *P. fluorescens* populations.

Here, we employ two global profiling approaches in *P. fluorescens* to identify RsmE- associated secretions that contribute to spatial structure formation beyond the already characterized extracellular polysaccharide and biosurfactant. To build on a previous study in *P. putida* that suggests indirect regulation by RsmE binding to other regulator mRNA (24), we first compare genes that are differentially expressed in Δ*rsmE* relative to the WT *P. fluorescens* Pf0-1 through RNA sequencing (RNA-seq). We next identify extracellular proteins that are unique to Δ*rsmE* compared to WT through LC-MS/MS. Lastly, we focus on genes that overlap in our transcriptomics and proteomics dataset to knockout four select secretion genes for competitive fitness measurements and microscopy analysis. Our systematic approach reveals RsmE- associated secretions that explain how Δ*rsmE* protects the spatial structure from the encroachment of neighboring cells and identifies additional candidates for future studies.

## RESULTS

### Diverse secretion genes are specifically upregulated in *ΔrsmE*

The primary goal of our study is to identify RsmE-associated extracellular secretions, beyond the previously characterized extracellular polysaccharide and biosurfactant, that contribute to the formation of dominant spatial structures in a crowded population (16). The Rsm/CsrA family of proteins are canonically known to function as a post transcriptional regulator that binds specific mRNAs to directly prevent their translation (25–27). However, recent studies have identified various mRNA-binding partners associated with transcription regulation (20–24), indicating that Rsm homologs could also indirectly regulate the production of certain secretions through altered transcription. We thus sought to identify secretion genes that are differentially expressed in *ΔrsmE* compared to WT through RNA-seq. As expected, the *rsmE* transcript (*Pfl01_1912*) was absent in the *ΔrsmE* dataset. Overall, 31 genes were observed to be upregulated and 17 were downregulated in *ΔrsmE* (Table 1 and Figure 1).

**Figure 1.**
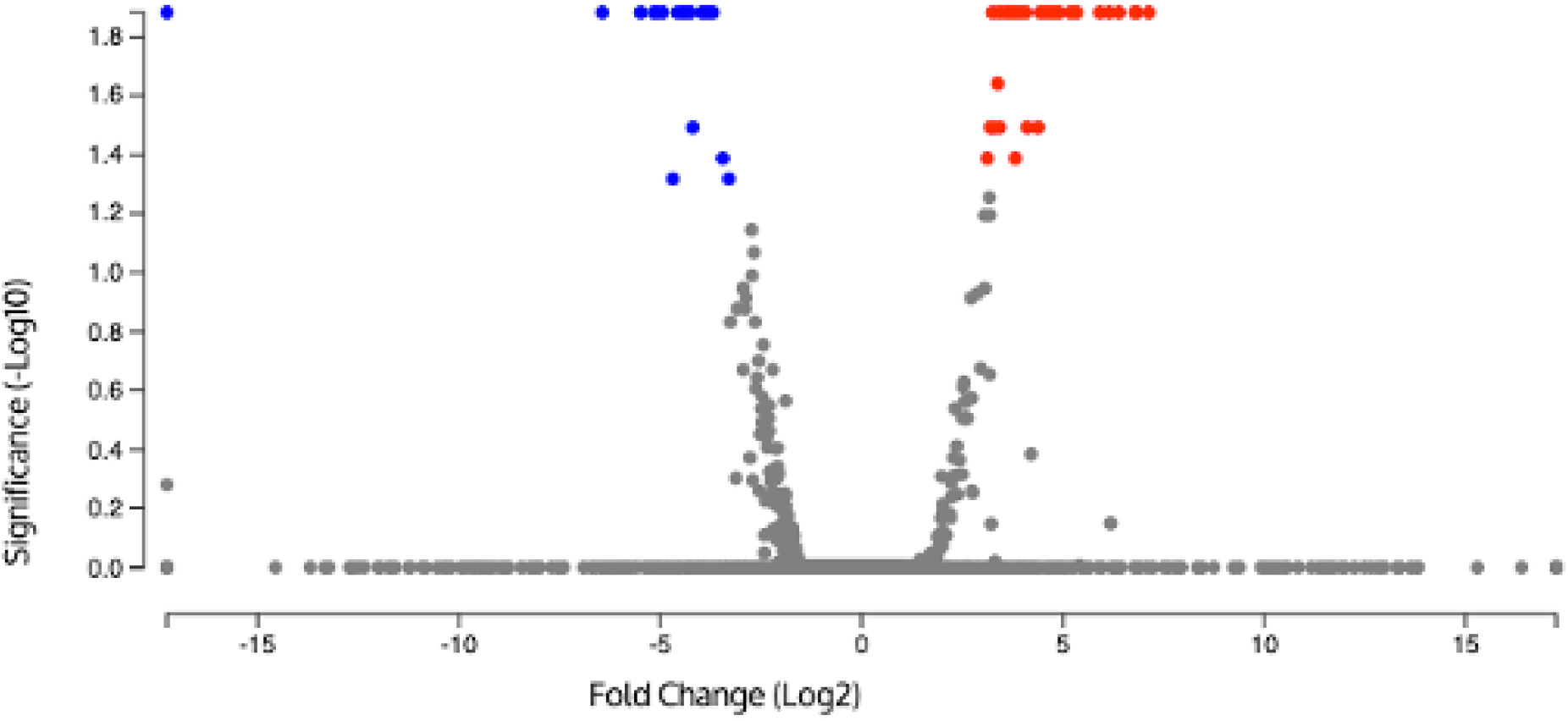
48 genes are differentially expressed in *ΔrsmE* relative to WT. Volcano plot comparing the relative expression of genes in *ΔrsmE* to WT as determined by RNA-seq. Genes with a fold change cutoff of 1-Log_2_ and FDR of 0.05 (shown as Log_10_) are labeled in blue for downregulation and red for upregulation, and genes with non-significant changes are labeled in grey. 31 genes are upregulated and 17 (excluding *rsmE*) are downregulated in *ΔrsmE*.

**Table 1.**
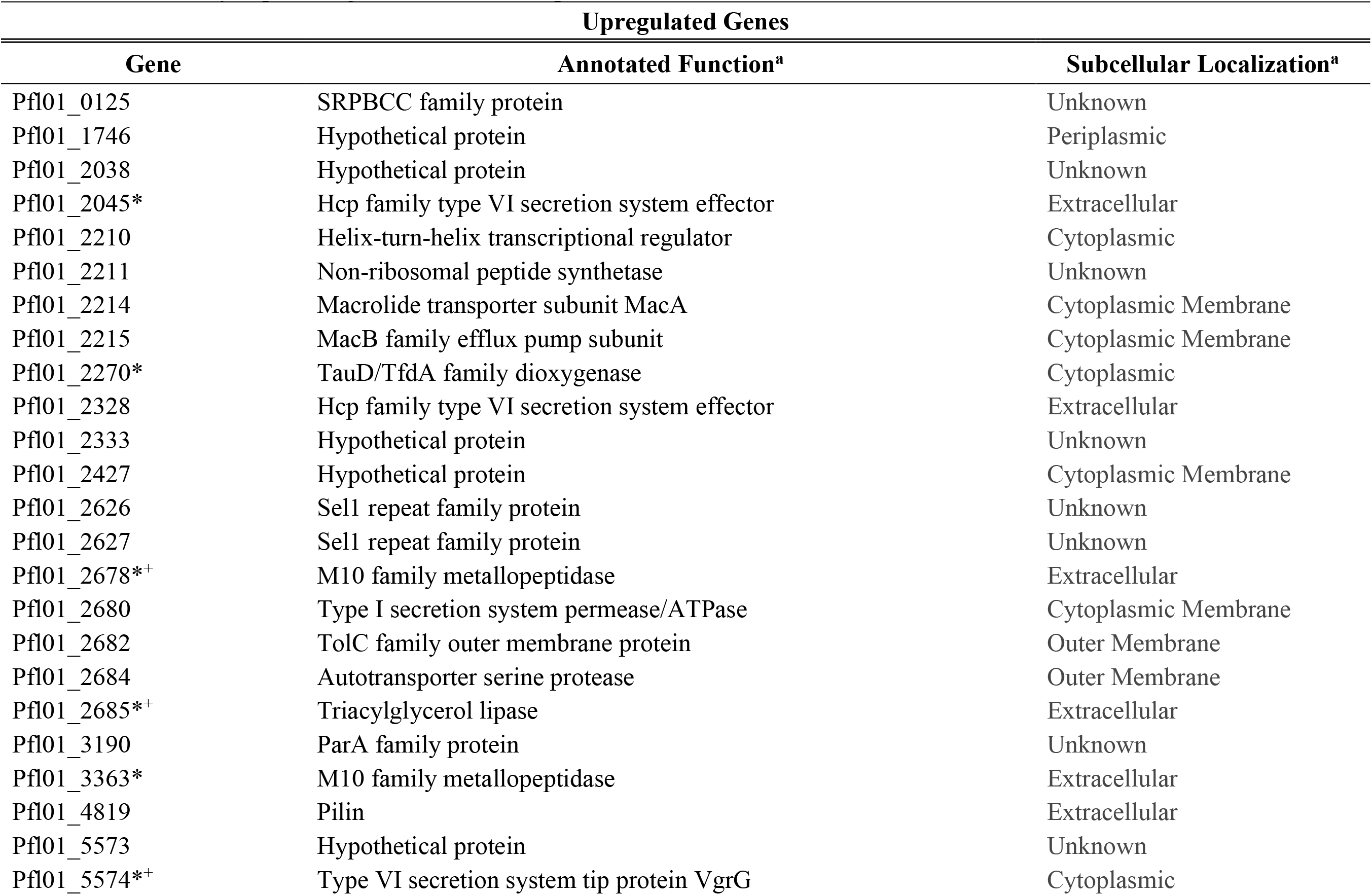
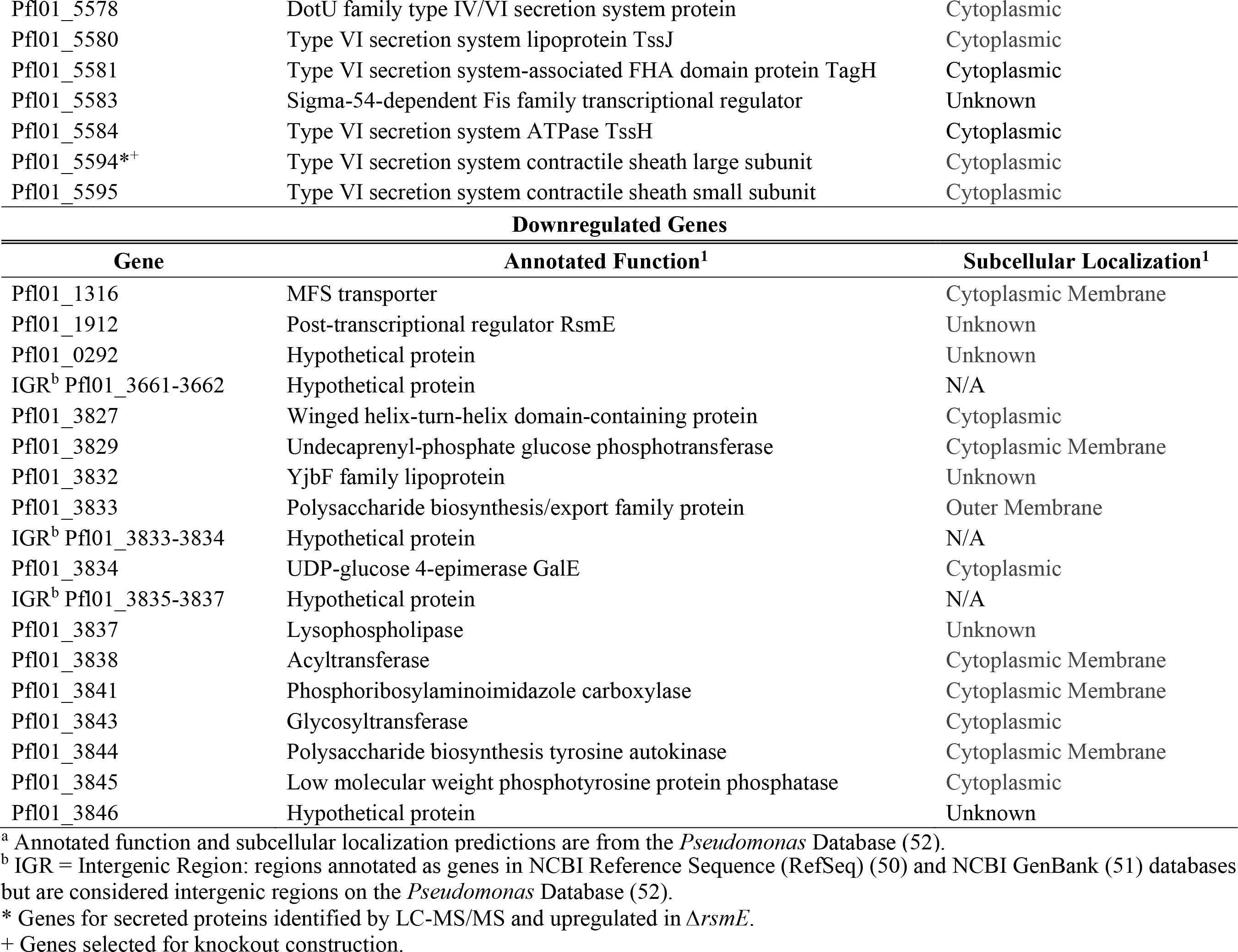
Differentially expressed genes in Δ*rsmE* compared to WT.

Aside from several hypothetical protein genes and a transporter-like gene, all downregulated genes clustered within the *Pfl01_3827*-*Pfl01_3846* region. The latter observation was unexpected, as we had previously described this gene cluster to produce the glucose-rich extracellular polysaccharide that drives the mucoid morphology of *ΔrsmE* (14) and space expansion within an overcrowded population (16). Since *ΔrsmE* produces significantly greater amounts of this extracellular polysaccharide compared to WT (14), indirectly upregulating the associated biosynthesis genes (*Pfl01_3827*-*Pfl01_3846*) via RsmE is clearly not the underlying mechanism. In contrast, *Pfl01_2211*, which encodes the biosynthesis gene for the biosurfactant that is uniquely produced in *ΔrsmE* and functions to locally sequester the extracellular polysaccharide (16), was upregulated in *ΔrsmE*, suggesting indirect regulation by RsmE. Immediately flanking genes (*Pfl01_2210*, *Pfl01_2214*, and *Pfl01_2215*) were also upregulated, that likely regulate and secrete the biosurfactant.

Other upregulated genes in *ΔrsmE* include diverse enzymes, transcription regulators, hypothetical proteins, and those associated with a type VI secretion system (T6SS) (Table 1). In particular, nearly a third of the genes in this dataset is predicted to encode the T6SS apparatus and its effectors. In addition, *Pfl01_5573* (hypothetical protein) and *Pfl01_5583* (transcription regulator) also fall within the T6SS gene cluster. The T6SS is best described as a bacterial weapon utilized in ecological warfare, comprising a spear-like apparatus loaded with diverse degradative effector proteins (e.g. proteases, nucleases, and lipases) which fires from the cytoplasm of the attacker to physically stab and deliver the effectors into the nearby competitors (28, 29). Unsurprisingly, producers of the T6SS also produce appropriate neutralizing anti- effector molecules for protection. Additional candidates that could contribute to RsmE- dependent spatial structure formation include upregulated gene products that are predicted to be secreted (Table 1).

### Knocking out *rsmE* dramatically increases the presence of extracellular proteins

Although we have identified upregulated genes in *ΔrsmE*, there is an obvious functional gap toward identifying those that contribute to spatial structure formation. We thus sought to identify extracellular proteins that are uniquely produced in *ΔrsmE* by comparing the extracellular proteomes of *ΔrsmE* and WT through LC-MS/MS. WT generated a total of 36,559 MS spectra reads mapping to 191 proteins in the genome, and *ΔrsmE* produced 36,138 MS spectra reads mapping to 310 proteins (Table S3). All 191 proteins found in WT were also found in *ΔrsmE*. To define proteins unique to *ΔrsmE*, a strict empirical abundance threshold of exactly 0 spectrum counts in WT was applied, which yielded 226 proteins. To further control for potential false discoveries arising from shared peptides assigned by the software’s parsimony grouping (30), the identified proteins were subsequently sorted based on 0% protein identification probability in WT, revealing 119 proteins as completely exclusive to *ΔrsmE* (Figure 2A, Table S3).

**Figure 2.**
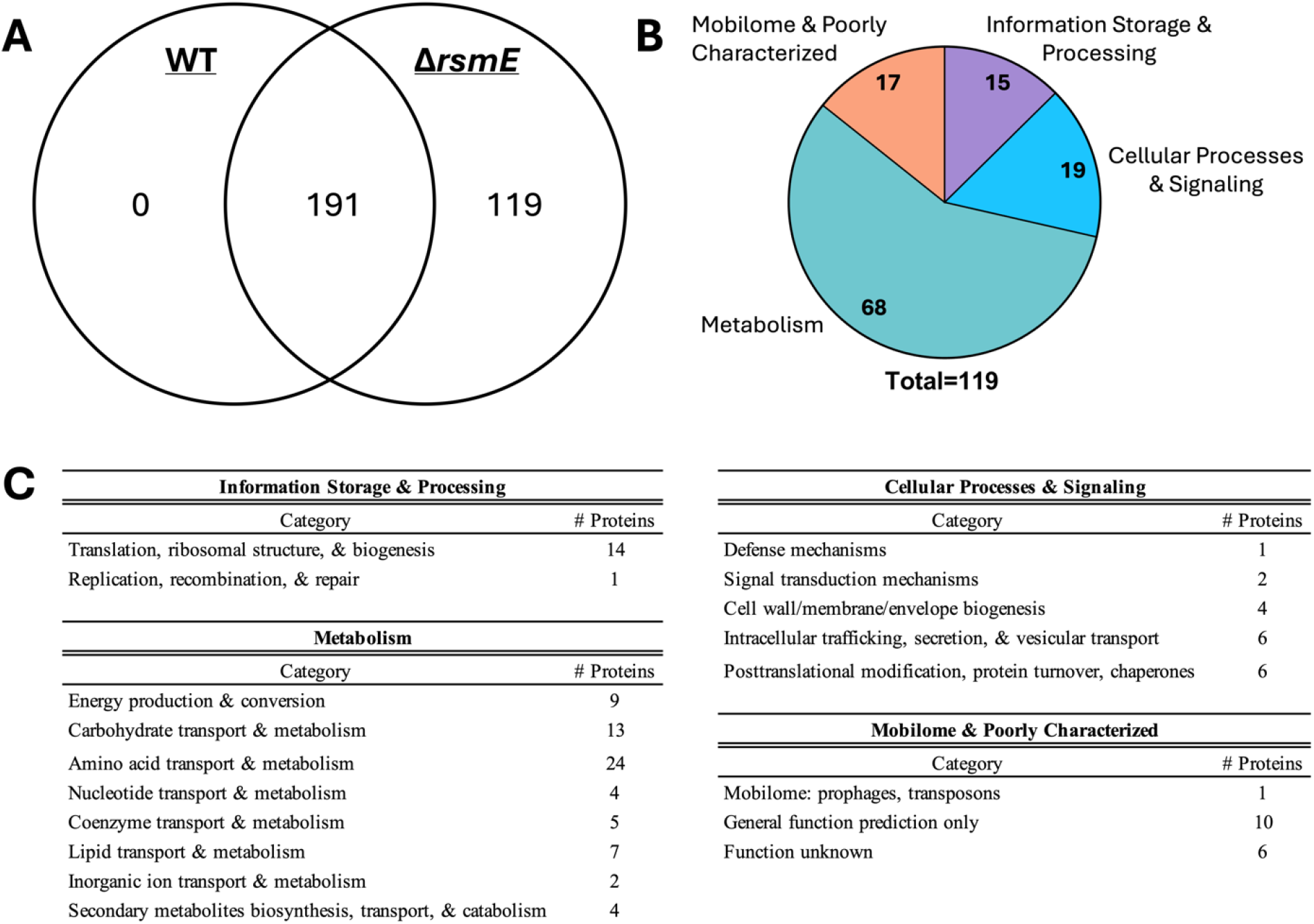
Absence of RsmE dramatically alters the extracellular proteome. (A) Venn diagram summarizing the 310 extracellular proteins identified using LC-MS/MS. Of the identified proteins, 191 are common to both WT and *ΔrsmE*, while 119 are unique to *ΔrsmE*. (B) The 119 proteins unique to *ΔrsmE* were classified into the four main categories of the Clusters of Orthologous Genes (COG) database (31). (C) Proteins were further categorized into the 26 specific functional sub-categories of the COG database (31).

The proteins unique to *ΔrsmE* were first sorted into 4 main categories using the Clusters of Orthologous Genes (COG) database (Figure 2B), then further categorized into the 26 specific functional categories of the COG database (Figure 2C, Table S4) (31). Nearly 60% of the *ΔrsmE*-specific proteins are involved in metabolism, including energy production/conversion or transport/metabolism of amino acids, carbohydrates, nucleotides, lipids, inorganic ions, and secondary metabolites. 16% of the proteins are involved in cellular processes and signaling, including signal transduction, posttranslational modification, secretion, and vesicular transport. 14% of the proteins are predicted to be hypothetical proteins or have annotations that are too general to be accurately categorized. Many of these proteins are predicted to be cytoplasmic or membrane associated, hence they likely stem from lysed cells during sample preparation. There is also increasing awareness that cytoplasmic and membrane proteins serve moonlighting functions outside the cell (32). For example, the function of elongation factor Tu is best known as a component of the translational machinery in the cytoplasm, but its moonlighting function in the extracellular space as a mediator of diverse matrix interactions is also acknowledged (33). Nevertheless, the fact that these proteins are exclusively detected in the *ΔrsmE* preparation provides a crucial path toward identifying RsmE-associated secretions that contribute to spatial structure formation.

### The T6SS confers a fitness advantage to *ΔrsmE* in competition with WT

The overlap across our transcriptomic and proteomic datasets effectively provides a short list of RsmE-associated candidates for functional analysis via engineering knockouts in *ΔrsmE*. None of the identified proteins match the down regulated genes in *ΔrsmE*. Among the proteins unique to *ΔrsmE*, four overlap with the upregulated genes in our RNA-seq dataset for *ΔrsmE* (Tables 1 and S3): T6SS sheath protein, T6SS tip protein, and two predicted extracellular peptidases. There are three additional proteins that overlap with the RNA-seq dataset and are enriched in *ΔrsmE* compared to WT: T6SS effector protein (0 spectrum counts and 16% protein probability in WT), predicted extracellular lipase (0 spectrum counts and 13% protein probability in WT), and predicted cytoplasmic oxygenase (1 spectrum count and 67% protein probability in WT). Due to the significant proportion of the candidates being associated with the T6SS, we selected two of the respective structural genes, *Pfl01_5574* and *Pfl01_5594*. We also selected two genes associated with abundantly produced enzymes in *ΔrsmE*: *Pfl01_2678*, encoding a putative extracellular peptidase, and *Pfl01_2685*, encoding a putative extracellular lipase. Each engineered knockout isolate produced the same colony morphology as the parent *ΔrsmE*, and neither extracellular polysaccharide (Figure S1A) nor biosurfactant (Figure S1B) production was affected by the respective gene deletion.

Although Δ*rsmE* and WT exhibit the same growth rate in both liquid and colony pure cultures, Δ*rsmE* spatially dominates the WT in co-culture colonies (12, 16). If the four selected genes contribute to the dominance of Δ*rsmE*, knocking them out in Δ*rsmE* should then reduce the relative fitness over the WT. We thus competed each secretion knockout mutant against the WT and compared their relative fitness to Δ*rsmE* (Figure 3A). As previously observed (16), Δ*rsmE* outcompetes the WT, as indicated by a relative fitness value greater than one. When *Pfl01_2678* and *Pfl01_2685* are knocked out in the Δ*rsmE* background, the relative fitness over WT does not significantly differ from that of Δ*rsmE*, indicating that the respective enzymes do not measurably contribute to spatial competition. However, when the T6SS genes (*Pfl01_5574* and *Pfl01_5594*) are individually knocked out in Δ*rsmE*, the resulting mutants both showed a significant decrease in the relative fitness over the WT compared to Δ*rsmE*. This reduction in relative fitness was comparable to that of M^S^*, an isolate in which *rsmE* is nonfunctional due to a frameshift deletion (denoted as M) and two previously identified RsmE-associated secretion genes (extracellular polysaccharide *Pfl01_3834* (denoted as *) and biosurfactant *Pfl01_2211* (denoted as ^S^)) are both deleted (16). We had previously demonstrated that M and Δ*rsmE* are phenotypically identical (12). These results indicate that the T6SS contributes to Δ*rsmE*’s spatial advantage over the WT, like the extracellular polysaccharide and biosurfactant.

**Figure 3.**
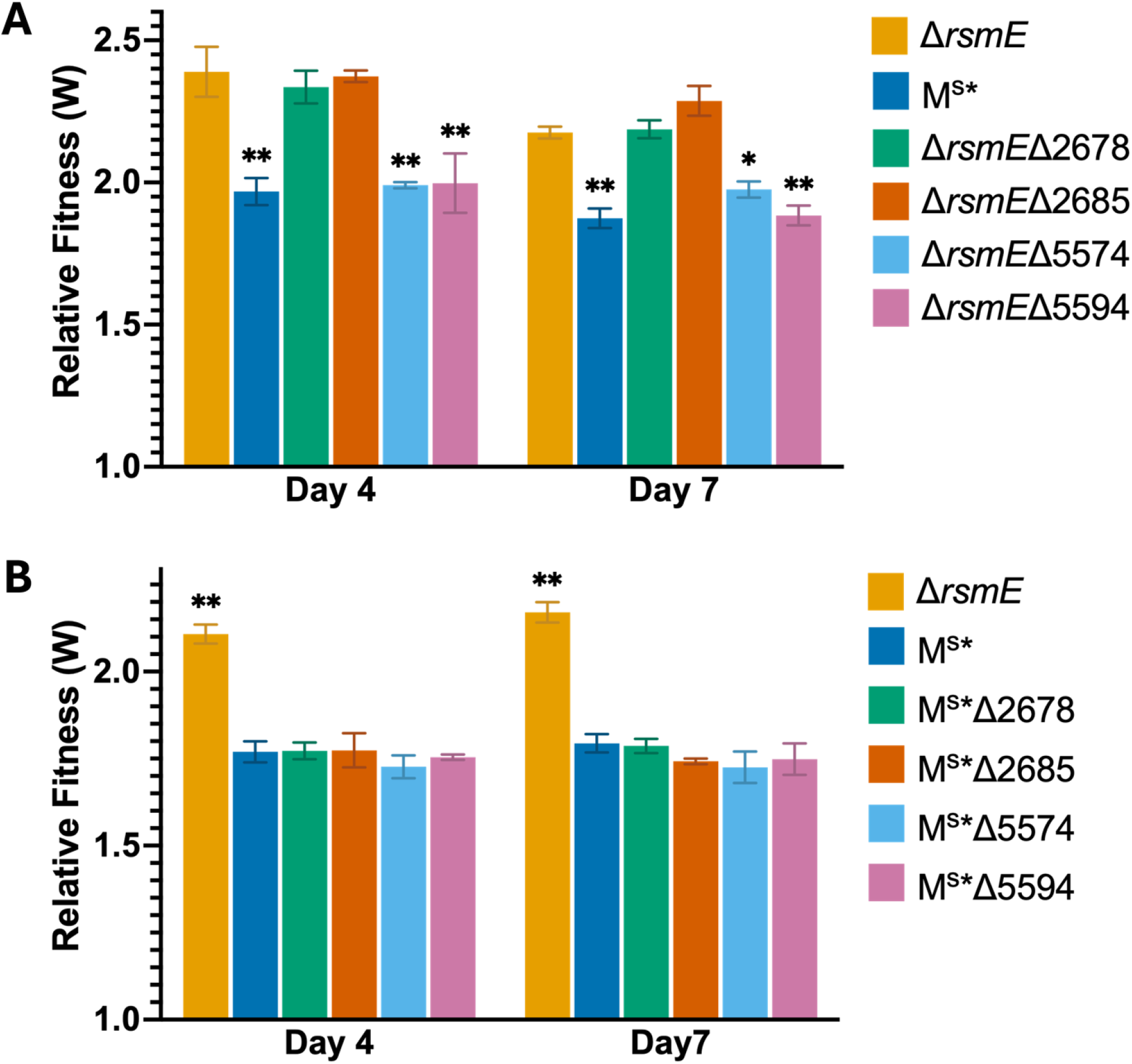
T6SS contributes to the dominance of *ΔrsmE* over the WT but not that of M^S^*. Relative fitness of candidate gene knockouts in (A) *ΔrsmE* and (B) M^S^* versus WT. Mutants were chromosomally tagged with kanamycin resistance, and WT was chromosomally tagged with streptomycin resistance. Mutant strains were competed with WT at a ratio of 10^-3^:1 on PAF, and three independent competitions were sampled after four and seven days of incubation. Error bars represent the standard deviations of the mean relative fitness of the mutant over WT (n=3). A relative fitness value (W) of 1 indicates the mutant is equally fit to WT, and W > 1 indicates that the mutant outcompeted the WT. Data at each time point were analyzed by two-way ANOVA (*P* < 0.0001), and Tukey’s honest significant different test was used to determine significance of all pairwise comparisons. Significance, denoted by * (*P* < 0.01) and ** (*P* < 0.0001), are noted compared to the *ΔrsmE*-WT control competition in (A) and compared to the M^S^*-WT control competition in (B).

To assess the combinational effect of the four candidate genes with the extracellular polysaccharide and biosurfactant, they were independently knocked out in M^S^*. When each knockout isolate was competed against the WT, none showed a further decrease in the relative fitness compared to their parent strain, M^S^* (Figure 3B). Thus, there appears to be no additive effect of losing the T6SS or the two enzymes when both the extracellular polysaccharide and biosurfactant are absent. This observation contrasts from the additive effect of the extracellular polysaccharide and biosurfactant in Δ*rsmE* (16). Furthermore, additional RsmE-associated products contribute to Δ*rsmE*’s fitness advantage over the WT, since T6SS or the two enzyme knockouts in M^S^* still outcompete the WT.

### T6SS kills WT cells that infiltrate *ΔrsmE*’s spatial structure

Diverse loss-of-function mutations in *rsmE* naturally emerge in WT colonies as mucoid patches, individually stemming from a single *rsmE* mutant cell that spatially dominates the neighboring WT cells over time (12). This process could be recapitulated and visualized with confocal microscopy by mixing fluorescently tagged WT cells (10^6^-10^7^ CFU) and a low number (10-100 CFU) of the isolated *rsmE* mutants or engineered *ΔrsmE* cells (12, 16). To assess the spatial structure formation of the gene knockouts, underrepresented GFP-labeled mutants were mixed with dsRedExpress-labeled WT and emergent patches were imaged, as described previously (12, 16). However, no obvious changes in spatial structure formation were observed (Figure S2). To better assess the role of the T6SS in spatial structure formation, unlabeled *ΔrsmE* and its T6SS mutants were mixed at a low relative frequency to GFP-labeled WT in the presence of propidium iodide and emerging non-fluorescent patches were visualized. A fluorescent signal in the red channel indicates the accumulation of propidium iodide binding to intracellular DNA within dead cells and/or extracellular DNA (34). At 20x optical magnification, we observed significantly greater red signal inside the patches formed by *ΔrsmE* and its peptidase or lipase knockouts compared to the T6SS mutants (Figure 4A). In contrast, we detected little to no red signal or intact cells at 100x optical magnification. To test the hypothesis that the red signal observed at 20x magnification likely represents extracellular DNA (eDNA), we examined extracted DNA under the same experimental settings. Indeed, propidium iodide binding to eDNA was not visible at 100x magnification but visible at lower magnifications (Figure S3). Given that the red signal is significantly reduced only within the patches formed by the T6SS mutants, it likely represents eDNA that is released from killed cells. However, it is unclear whether the eDNA originates from *ΔrsmE* or WT cells.

**Figure 4.**
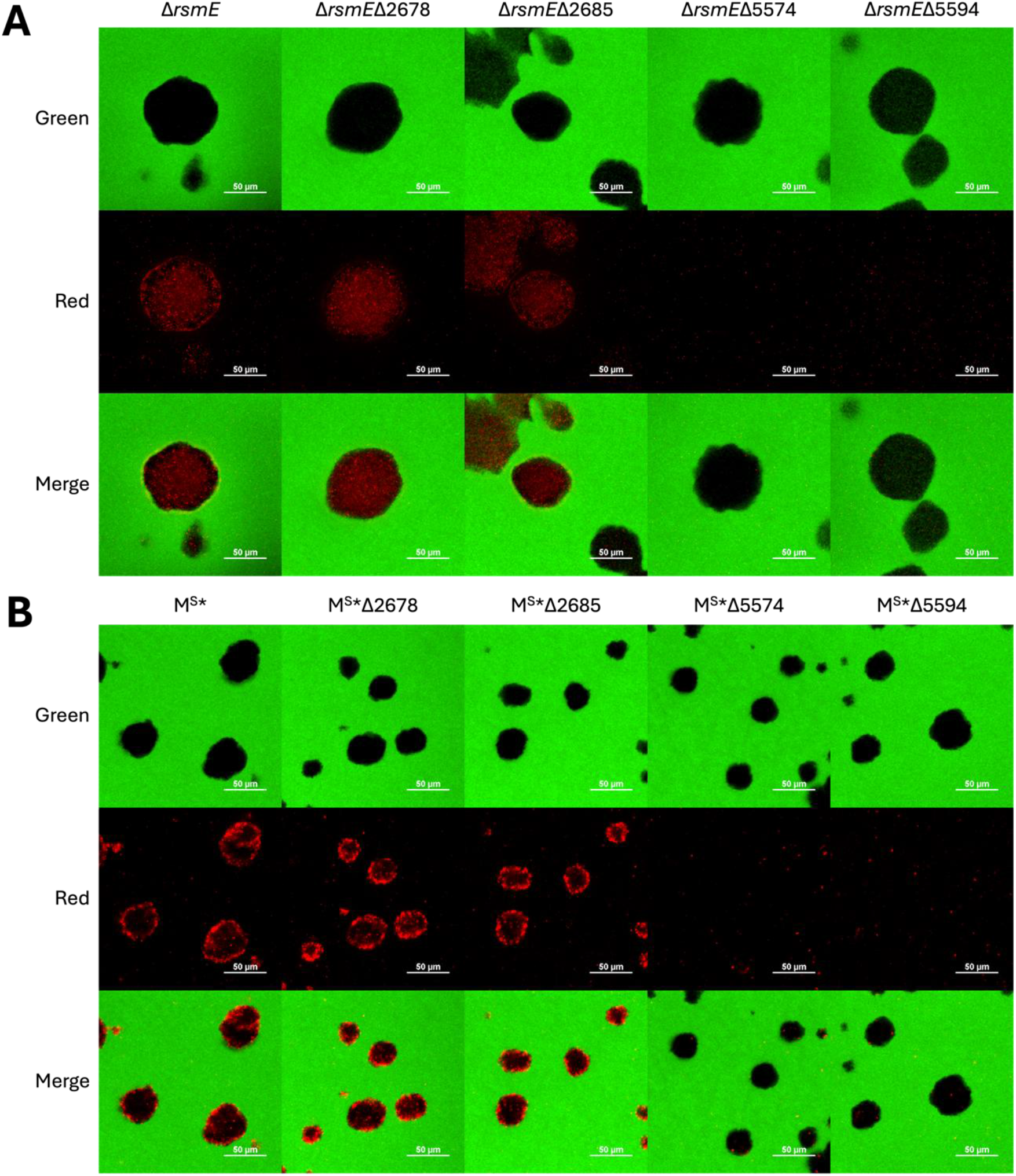
T6SS kills WT cells within patches formed by *ΔrsmE*. Unlabeled mutants, as noted above each panel, were mixed with GFP-labeled WT at a 10^-5^ relative frequency and incubated for three days with propidium iodide supplemented to each initial inoculum. Confocal images of the emergent patches were generated using the 20x PLAN APO objective using the green, red, and merge channels. (A) DNA released from dead cells (red) are seen in high abundance throughout the patches formed by Δ*rsmE*, Δr*smE*Δ2678 (metalloprotease), and Δ*rsmE*Δ2685 (triglycerol lipase). In contrast, death is rarely associated with the patches formed by Δ*rsmE*Δ5574 and Δ*rsmE*Δ5594, that lack the T6SS. (B) PI signal is concentrated at the boundary of the patches formed by M^S^*, M^S^*Δ2678, and M^S^*Δ2685 where the mutant cells contact WT cells. Similar to the T6SS knockouts in the Δ*rsmE* background, M^S^*Δ5574 and M^S^*Δ5594 do not show cell death. Scale bars represent 50 µm.

A clear difference in the spatial structure formed by M^S^* compared to that of *ΔrsmE* in WT colonies is the absence of cell-free space (i.e. low cell density) and reduced patch size, wherein the M^S^* cells are tightly packed (Figure S2) (16). If the red signal detected within *ΔrsmE* patches represent eDNA from *ΔrsmE* cells, then the same signal should be observed uniformly throughout M^S^* patches in the presence of propidium iodide. However, the red signal is specifically intensified at the border of M^S^* patches (Figure 4B). The same spatial pattern is produced by patches formed by M^S^* with either the peptidase or lipase knocked out, but it is abolished when the T6SS genes are knocked out in M^S^* (Figure 4B). These results collectively indicate that the eDNA is primarily stemming from the immediately surrounding WT cells.

Two hallmark features of the spatial structure formed by *ΔrsmE* within WT colonies are dramatically reduced cellular density and prevention of WT encroachment (12, 16). However, our propidium iodide experiment (Figure 4A) indicates that the neighboring WT cells may not be physically blocked from infiltrating *ΔrsmE*’s genotypic boundary. Indeed, visualizing patches formed by unlabeled *ΔrsmE* T6SS mutants in a GFP-labeled WT colony, revealed the presence of WT cells at 100x optical magnification (Figure 5A). In contrast, WT cells were essentially absent within patches of M^S^*, *ΔrsmE*, and the two enzyme knockouts (Figure 5A). When the fluorescence intensity from encroaching GFP-labeled WT cells was quantified, there is a significant increase in fluorescence signal within the mutant patch when the T6SS is deleted in the *ΔrsmE* background but not the M^S^* background (Figure 5B). The unique presence of WT cells within the *ΔrsmE* T6SS patches also coincides with the unique detection of eDNA (Figure 4A). Taken together, these observations strongly suggest that some WT cells do encroach into *ΔrsmE*’s space, but they are effectively killed by the T6SS. However, the functional roles of the putative peptidase and lipase in spatial structure formation, if any, remain unclear.

**Figure 5.**
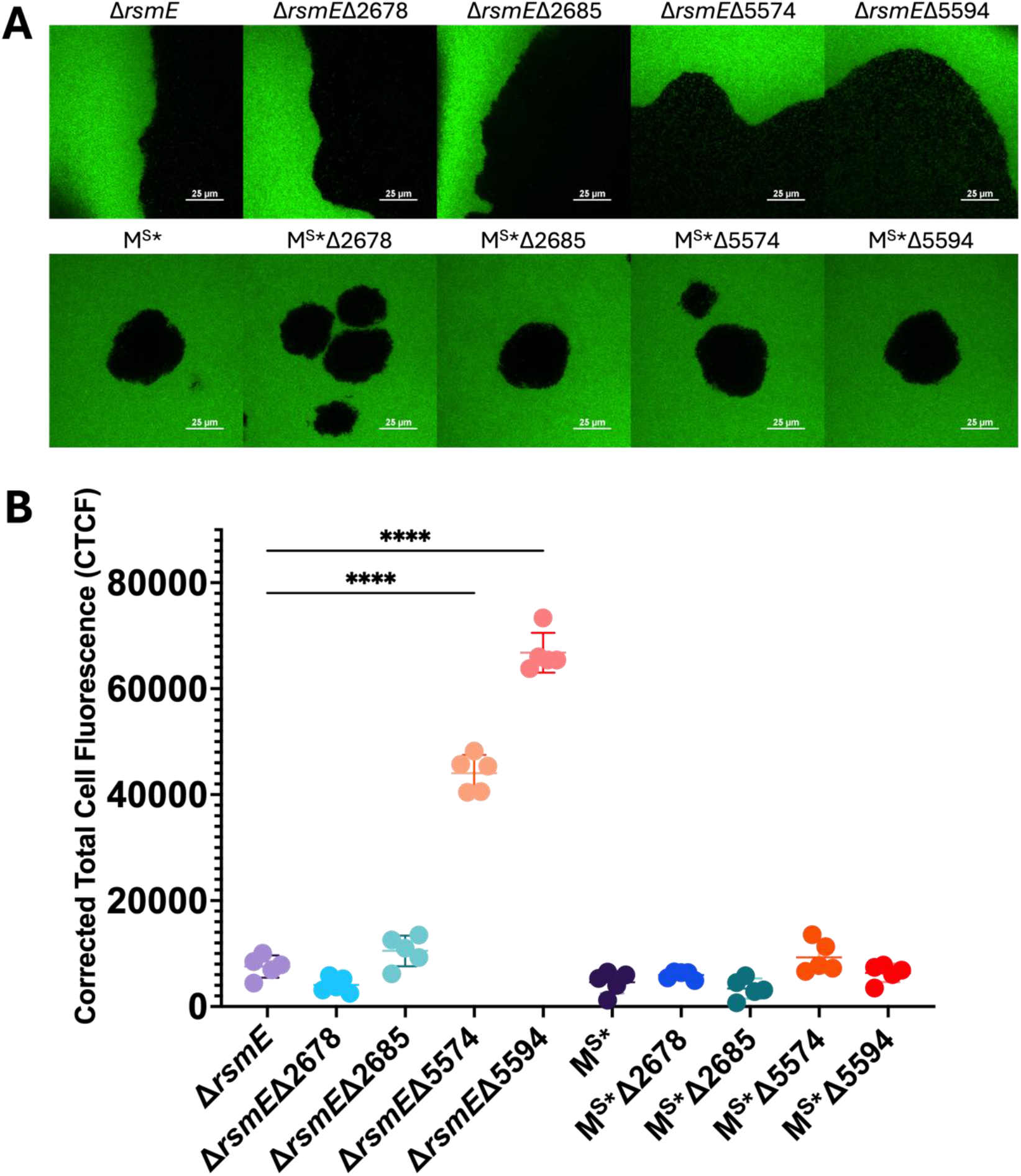
T6SS prevents the encroachment of WT cells into the Δ*rsmE* patch. (A) Unlabeled mutants, as noted above each panel, were diluted to a 10^-5^ relative frequency, mixed with undiluted GFP-labeled WT, and imaged after three days of growth. Confocal images were captured using the 100x TU PLAN APO objective. Only the T6SS knockouts in the Δr*smE* background (Δ*rsmE*Δ5574 and Δ*rsmE*Δ5594), have green WT cells within the patches, suggesting that the T6SS is responsible for killing encroaching WT cells. None of the knockouts in M^S^* show encroachment of WT cells, as M^S^* patches are tightly packed with only M^S^* cells. Scale bars represent 25 µm. (B) Corrected Total Cell Fluorescence (CTCF) of encroaching GFP- labeled WT cells was measured in five areas within each mutant patch. Deletion of the T6SS in Δr*smE* significantly increases the measured fluorescence within the patch. However, deletion of the T6SS in M^S^* does not significantly increase fluorescence intensity. Data were analyzed by ANOVA (*P* < 0.0001) and Tukey’s honest significant different test (*P* < 0.0001). Significance (****) is shown compared to *ΔrsmE*.

## DISCUSSION

The two global profiling approaches utilized in this study collectively identified a list of candidate genes that are either directly or indirectly regulated by RsmE in *P. fluorescens* Pf0-1. The canonical mechanism by which Rsm homologs function is by inhibiting the translation of bound mRNA (26). A recent study conducted in *P. putida* KT2440 showed that RsmE binds to the transcripts of 241 genes, including several regulators (24). Under this functional model, one would expect to observe changes primarily in the proteome in the absence of RsmE, but changes in the transcriptome should also be expected if RsmE alters the translation of other regulatory systems. Indeed, our LC-MS/MS approach identified 119 uniquely present proteins in extracellular preparations of *ΔrsmE*, while our RNA-seq approach revealed a relatively modest number of 48 differentially expressed genes. Furthermore, the bulk of these genes are predicted to encode proteins that participate in the biosynthesis and transport of common extracellular secretions, including the extracellular polysaccharide (14) and biosurfactant (16), in addition to the T6SS and its putative effectors identified in this study. Although our proteomics data support the functional role of RsmE as a major regulator of extracellular secretions, the key elements that drive the formation of spatially dominant structures appear to be indirectly modulated by RsmE.

RsmE-modulated extracellular polysaccharide and biosurfactant function together to create and maintain structured space of low cell density in a crowded colony. The polysaccharide is solely responsible for creating the space by pushing away the neighboring cells, and the biosurfactant appears to locally sequester other extracellular secretions to physically maintain the genotypic boundary (12, 16). Our study demonstrates that RsmE also modulates the production of a T6SS, which likely kills neighboring cells to protect the spatiogenetic structure. The T6SS is a well-known bacterial weapon in ecological warfare, and RsmA of *P. aeruginosa*, which lacks RsmE, has been demonstrated to regulate its production (35). The tip of the T6SS, which physically punctures through the membrane of a neighboring cell, is loaded with diverse degradative effector proteins that function to kill the competitor (28, 36). The T6SS likely plays an important role in self vs non-self recognition (28).

The phenotypic noise associated with the production of the T6SS within an isogenic population (37–39) could result in occasional self-killing. However, we provide several layers of evidence that it is primarily the WT rather than *ΔrsmE* cells that are killed within the mutant patch: 1) *ΔrsmE* T6SS mutants exhibit significantly reduced relative fitness compared to *ΔrsmE* in competitions against the WT, 2) live WT cells are present within the *ΔrsmE* patch exclusively when the T6SS is knocked out, and 3) death is localized to the M^S^* patch boundary where the M^S^* and WT cells are in direct contact with one another. Since the M^S^* patch is tightly packed with M^S^* cells (16), a uniform pattern of death should have been observed throughout the patch and at the patch boundary if death is primarily caused through self-killing. In contrast, we observed a relatively uniform pattern of death within the *ΔrsmE* patch, which also coincides with the uniformly low cell-density both across the patch and its border (12, 16). Knocking out the T6SS in M^S^* did not measurably impact its relative fitness compared to M^S^* in competitions against WT, which is likely due to the fact that physical interactions between M^S^* and WT cells are spatially limited to the genotypic boundary. We would also expect to observe an increase in the relative fitness of the T6SS knockout in M^S^* if there was significant self-killing.

The producer of a T6SS also produces immunity proteins that neutralize the activity of each effector protein (28, 36). Therefore, a *ΔrsmE* cell is expected to be immune to the attack from another *ΔrsmE* cell, but a WT cell should succumb to *ΔrsmE*’s attack. There is increasing evidence that a T6SS attack could also be neutralized through non-specific means, including the production of protective extracellular polysaccharides to create a physical barrier or relying on the activation of envelope stress pathways as a general protective measure (36). Such immunity-independent mechanisms could be uniquely employed by *ΔrsmE*, particularly through multiple extracellular secretions that are lacking in WT. Along with killing neighboring cells via direct cell-cell contact, the T6SS in various species is also known to have indirect contact functions that can be beneficial in dense, competitive environments, such as acquisition of metal ions and resistance to oxidative stress (40–42).

Our analysis of the RsmE-modulated peptidase and lipase showed no measurable effect on spatial structure formation or relative fitness over the WT. Rather than being directly involved in spatial structure formation, they may be involved in nutrient uptake (43) or their benefit may be masked by other secretions (44). Much like the reduced influence of the T6SS in the absence of the extracellular polysaccharide and biosurfactant, realized benefit may be dependent on specific combinations of diverse secretions present in the extracellular matrix. Additional RsmE- modulated extracellular proteins identified in this study, including numerous hypothetical proteins, represent a rich resource for exploring unique mechanisms of social interaction in structured populations and characterizing the hierarchical regulation surrounding RsmE.

## METHODS

### Bacterial strains and culture conditions

All *P. fluorescens* strains used in this study are listed in Table S1. *Escherichia coli* Jm109 (Promega) was used for routine cloning, and *E. coli* strains S17-λ*pir* and HB101 were used for conjugations (45, 46). Liquid and solid LB (Fisher) were used for routine growth. Difco Pseudomonas Agar F (PAF; Fisher) was used for all assays, phenotypic screens, competitions, and microscopy. *Pseudomonas* minimum medium (PMM; 3.5 mM Potassium phosphate dibasic trihydrate, 2.2 mM potassium phosphate monobasic, 0.8 mM ammonium sulfate, 100 mM Magnesium sulfate, 100 mM sodium succinate) was used to selectively isolate *P. fluorescens* from conjugations with *E. coli*. Antibiotics were added to the media when needed at the following final concentrations: kanamycin (50 µg/mL), gentamicin (20 µg/mL), ampicillin (100 µg/mL), and streptomycin (50 µg/mL). *P. fluorescens* was cultured at 30 °C or ambient room temperatures (∼22 °C) as indicated, while *E. coli* was grown at 37 °C. Liquid cultures were shaken at 250 rpm while incubating.

### RNA purification and RNA-sequencing

For total RNA purification, WT and Δ*rsmE* were grown overnight in 5 mL cultures to saturation. Then, 20 µL of each culture was spotted in triplicate on PAF plates and grown at room temperature for 3 days. After incubation, the resulting colonies were harvested, and RNA was extracted from each colony sample using Trizol reagent (Invitrogen) using the manufacturer’s protocol. RNA was quantified with the NanoDrop One (Thermo Scientific) spectrophotometer and the A260/A280 ratio calculated to assess purity. RNA samples were sequenced at SeqCenter (Pittsburgh, PA).

Raw sequencing reads were processed using FastQC V0.11.5 (http://www.bioinformatics.babraham.ac.uk/projects/fastqc/) to assess the quality of each data set. Reads were then aligned with the *P. fluorescens* Pf0-1 genome using HISAT-2 v2.1.0 (47) and Bowtie2 V2.3.2 (48). Cufflinks v2.2.1 (49) was then used to assemble the transcripts, and Cuffdiff v2.2.1 (49) was used to identify differentially expressed genes with Log2 fold change of 1 and a false discovery rate adjusted p-value (FDR) of 0.05. Gene annotations are from the NCBI Reference Sequence (RefSeq) (50), NCBI GenBank (51) and *Pseudomonas* database (52), and subcellular localization information is from the *Pseudomonas* database (52).

### Identification of extracellular proteins

Twenty microliters of overnight cultures of WT and Δ*rsmE* were spotted in triplicate on PAF and incubated at room temperature for three days. After incubation, three entire colonies and the medium immediately surrounding the colony (∼20 mm^2^) were excised from the plate and placed in a 50 mL conical tube. Samples were immersed in protein extraction buffer (50 mM Tris, 150 mM NaCl, 1 mM PMSF, 1 mM EDTA, pH 7.6) and vortexed until the cells were separated from the agar slices. The buffer was then transferred to a new tube, leaving the agar slices behind. The contents of the tube were passed through a 0.2 µm filter and collected in a new tube. The filtered supernatant was concentrated using a 3 kDa MWCO PES concentrator tube, and 400 µg of each sample was loaded onto an SDS-PAGE gel and run at 100V until the entire sample entered into the stacking gel (4% acrylamide). The gel was then fixed with 25% acetic acid. Samples were excised then analyzed at Michigan State University Proteomics Core (East Lansing, MI) for LC-MS/MS analysis. LC-MS/MS data was analyzed with scaffold viewer (Proteome Software, Inc). Spectra reads were aligned with the Pf0-1 genome. Protein threshold was set at 1.0% FDR, and peptide threshold was set at 0.1% FDR with the minimal number of peptides of 3. Proteins were classified as unique to *ΔrsmE* if they contained 0 spectrum counts and a 0% protein probability in WT. Annotated function and subcellular localization predictions are from the *Pseudomonas* database (52), and proteins unique to Δ*rsmE* were functionally categorized using the Clusters of Orthologous Genes (COG) database (31).

### Construction of knockout mutants and chromosomal markers

Knockout mutants were generated as previously described using the gene splice by overlap extension method (16, 53). Briefly, for each targeted gene, ∼500bp directly upstream and downstream of the gene was amplified using the up set and down set of primers (Table S2) respectively and subsequently joined together by PCR using the forward up primer and the reverse down primer. The ∼1000 bp joined flanking regions were cloned into pGEMT-Easy, then subcloned into the suicide plasmid pMQ30 and transformed into *E. coli* S17.1λpir. Mating was then used to transform *ΔrsmE* and M^S^* with pMQ30 and followed by sucrose selection as previously described (16). Resulting colonies were screened using the outside primer set (Table S2) for the expected reduction in amplicon size. Gene deletions were further confirmed through whole genome sequencing at SeqCenter (Pittsburgh, PA). All knockouts were chromosomally tagged with kanamycin-resistance for quantifying competitions and GFP for microscopy using the mini-Tn7 chromosomal insertion system (54) as previously described (12, 14, 16). The kanamycin and streptomycin resistance markers used in the competitions were previously demonstrated to be neutral for measuring relative fitness in *P. fluorescens* Pf0-1 (12).

### Competition assay

The kanamycin-resistant knockout isolates were competed against the streptomycin-resistant WT isolate as described previously (12, 16). Briefly, overnight cultures were washed and resuspended in 1 mL PBS, and the kanamycin-resistant isolates were diluted in PBS to 10^-3^ then mixed with equal volumes of the undiluted streptomycin-resistant WT isolate. Twenty microliters of each competition mixture were spotted onto PAF in triplicate and incubated at 22°C. The initial population sizes were enumerated by serially diluting each competition mixture, plating onto PMM supplemented with either kanamycin or streptomycin, and counting the resulting colonies the following day. After four and seven days of incubation, the spotted colonies were resuspended in 5 mL PBS, serially diluted, plated onto PMM supplemented with either kanamycin or streptomycin, and colonies were counted, just like the initial competition mixture. Relative fitness (*W*) of each knockout isolate against the WT was calculated as follows (55): [ln(CFU of mutant at time of sampling/CFU of mutant at time zero)]/[ln(CFU of WT at time of sampling/CFU of WT at time zero)].

### Detection of extracellular polysaccharide and biosurfactant production

To determine whether the mutants produce the extracellular polysaccharide, 20 μL of overnight culture was spotted onto PAF plates and imaged after one day of growth at room temperature. Extracellular polysaccharide production results in a mucoid colony morphology (12, 16). Biosurfactant production was determined using a Nucleopore Track-Etch polycarbonate membrane (Whatman; 0.4 μM pore size, 90-mm diameter). As previously described (12, 16), the membrane has a dull and shiny side due to the manufacturing process that allows for detection of biosurfactant production. The dull side contains gaps and ridges that trap cells but let the biosurfactant permeate through, producing a visible ring around the colony. A membrane with the dull side facing up was overlain on PAF plates using sterile forceps, and 20 μL of overnight culture was spotted on the membrane, allowed to dry, and imaged after one day of growth at room temperature. The presence of a ring around the colony indicates biosurfactant production. Images were captured using the Hayear overhead microscope (HY-2307).

### Microscopy

Overnight cultures of WT (either GFP- or dsRedExpress-labeled) and each mutant (either GFP- or non-labeled) were washed and resuspended in PBS. All mutant cell suspensions were serially diluted to 10^-5^ in PBS and mixed with equal volumes of the undiluted WT suspension. To visualize the effects of the T6SS, propidium iodide (Invitrogen) was used to detect cellular death under the assumption that a fluorescent signal in the red channel represents the accumulation of propidium iodide binding to intracellular DNA within dead cells and/or extracellular DNA. Competitions using propidium iodide were washed with 10 mM magnesium sulfate before resuspension in PBS, and propidium iodide was added to the inoculum at a concentration of 30 µg/mL. Twenty microliters of each competition mixture were spotted in triplicate on PAF plates and incubated at room temperature. Confocal microscopy was conducted as previously described, using an agar slice containing one entire colony that was placed on a microscope slide and visualized without a coverslip (16). Confocal microscopy was conducted using the Nikon Ti2 microscope with 10x Plan Apo, 20x Plan Apo, or 100x TU Plan Apo objective and the NIS Elements software. Images were rendered using the NIS Elements and Fiji (ImageJ) software.

### Fluorescence intensity quantification

Fluorescence intensity of encroaching cells was quantified using Fiji (ImageJ) software. For each mutant, five technical replicates were measured for each sample using an area of 10,000 square pixels in five separate positions within the mutant patch. The Corrected Total Cell Fluorescence (CTFC) for each sample replicate (n=5) was calculated using the raw integrated density values to avoid scaling artifacts and adjusted to background signal using the following formula: CTFC = Integrated Density – (Area of Selected Region * Mean Fluorescence of Background).

### Statistical analysis

The same statistical analyses were performed on both the relative fitness data and the fluorescence intensity data. Relative fitness data were collected for three biological replicates and two technical replicates per biological replicate. Fluorescence intensity of encroaching cells was collected for five replicates using five regions with a fixed area within the confocal image. To determine significance, the data were analyzed with a one-way analysis of variance (ANOVA) for fluorescence intensity and a two-way ANOVA for relative fitness, then Tukey’s Honest Significant Difference test (*P* < 0.05) was used to make pairwise comparisons within each dataset. Comparisons deemed significantly different are noted. Statistical tests were conducted using GraphPad Prism.

## Data availability

All noncommercial plasmids or strains used in this study are available for distribution upon request.

## ACKNOWLEDGEMENTS

W.K. designed the study; M.W., A.E., S.S., A.D., R.S., C.G., A.V., and W.K. performed experiments; M.W., A.E., S.S., and W.K. analyzed data; and M.W., A.E., S.S., and W.K. wrote the manuscript. This study was funded by the National Institute of General Medical Sciences of the NIH (1R15GM132856 to W.K.). Confocal imaging was supported with a grant from the NSF, DBI #1726368.

---

**Table S1.** *Pseudomonas* isolates used in this study.

| Isolate | Genotype | Source |
| --- | --- | --- |
| Pf0-1 | Wildtype (WT) | (56) |
| Pf0-1 GFP | WT (Tn7-Gfpmut2) | (12) |
| Pf0-1 RFP | WT (Tn7-dsRedExpress) | (12) |
| Pf0-1 SM | WT (Tn7-SmR) | (12) |
| $\Delta rsmE$ | WT ($\Delta Pfl01\_1912$) | (12) |
| $\Delta rsmE$ Km | WT ($\Delta Pfl01\_1912$, Tn7-KmR) | (12) |
| $\Delta rsmE$ GFP | $\Delta rsmE$ (Tn7-Gfpmut2) | (12) |
| M<sup>S*</sup> | WT (*rsmE*ΔA126, $\Delta Pfl01\_2211$, $\Delta Pfl01\_3834$) | (16) |
| M<sup>S*</sup> Km | WT (*rsmE*ΔA126, $\Delta Pfl01\_2211$, $\Delta Pfl01\_3834$, Tn7-KmR) | (16) |
| M<sup>S*</sup> GFP | M<sup>S*</sup> (Tn7-Gfpmut2) | (16) |
| $\Delta rsmE\Delta 2678$ | $\Delta rsmE$ ($\Delta Pfl01\_2678$) | This Study |
| $\Delta rsmE\Delta 2685$ | $\Delta rsmE$ ($\Delta Pfl01\_2685$) | This Study |
| $\Delta rsmE\Delta 5574$ | $\Delta rsmE$ ($\Delta Pfl01\_5574$) | This Study |
| $\Delta rsmE\Delta 5594$ | M<sup>S*</sup> ($\Delta Pfl01\_5594$) | This Study |
| M<sup>S*</sup>$\Delta 2678$ | M<sup>S*</sup> ($\Delta Pfl01\_2678$) | This Study |
| M<sup>S*</sup>$\Delta 2685$ | M<sup>S*</sup> ($\Delta Pfl01\_2685$) | This Study |
| M<sup>S*</sup>$\Delta 5574$ | M<sup>S*</sup> ($\Delta Pfl01\_5574$) | This Study |
| M<sup>S*</sup>$\Delta 5594$ | M<sup>S*</sup> ($\Delta Pfl01\_5594$) | This Study |
| $\Delta rsmE\Delta 2678$ Km | $\Delta rsmE$ ($\Delta Pfl01\_2678$, Tn7-KmR) | This Study |
| $\Delta rsmE\Delta 2685$ Km | $\Delta rsmE$ ($\Delta Pfl01\_2685$, Tn7-KmR) | This Study |
| $\Delta rsmE\Delta 5574$ Km | $\Delta rsmE$ ($\Delta Pfl01\_5574$, Tn7-KmR) | This Study |
| $\Delta rsmE\Delta 5594$ Km | M<sup>S*</sup> ($\Delta Pfl01\_5594$, Tn7-KmR) | This Study |
| M<sup>S*</sup>$\Delta 2678$ Km | M<sup>S*</sup> ($\Delta Pfl01\_2678$, Tn7-KmR) | This Study |
| M<sup>S*</sup>$\Delta 2685$ Km | M<sup>S*</sup> ($\Delta Pfl01\_2685$, Tn7-KmR) | This Study |
| M<sup>S*</sup>$\Delta 5574$ Km | M<sup>S*</sup> ($\Delta Pfl01\_5574$, Tn7-KmR) | This Study |
| M<sup>S*</sup>$\Delta 5594$ Km | M<sup>S*</sup> ($\Delta Pfl01\_5594$, Tn7-KmR) | This Study |
| $\Delta rsmE\Delta 2678$ GFP | $\Delta rsmE$ ($\Delta Pfl01\_2678$, Tn7-Gfpmut2) | This Study |
| $\Delta rsmE\Delta 2685$ GFP | $\Delta rsmE$ ($\Delta Pfl01\_2685$, Tn7-Gfpmut2) | This Study |
| $\Delta rsmE\Delta 5574$ GFP | $\Delta rsmE$ ($\Delta Pfl01\_5574$, Tn7-Gfpmut2) | This Study |
| $\Delta rsmE\Delta 5594$ GFP | M<sup>S*</sup> ($\Delta Pfl01\_5594$, Tn7-Gfpmut2) | This Study |
| M<sup>S*</sup>$\Delta 2678$ GFP | M<sup>S*</sup> ($\Delta Pfl01\_2678$, Tn7-Gfpmut2) | This Study |
| M<sup>S*</sup>$\Delta 2685$ GFP | M<sup>S*</sup> ($\Delta Pfl01\_2685$, Tn7-Gfpmut2) | This Study |
| M<sup>S*</sup>$\Delta 5574$ GFP | M<sup>S*</sup> ($\Delta Pfl01\_5574$, Tn7-Gfpmut2) | This Study |
| M<sup>S*</sup>$\Delta 5594$ GFP | M<sup>S*</sup> ($\Delta Pfl01\_5594$, Tn7-Gfpmut2) | This Study |

**Table S2.**
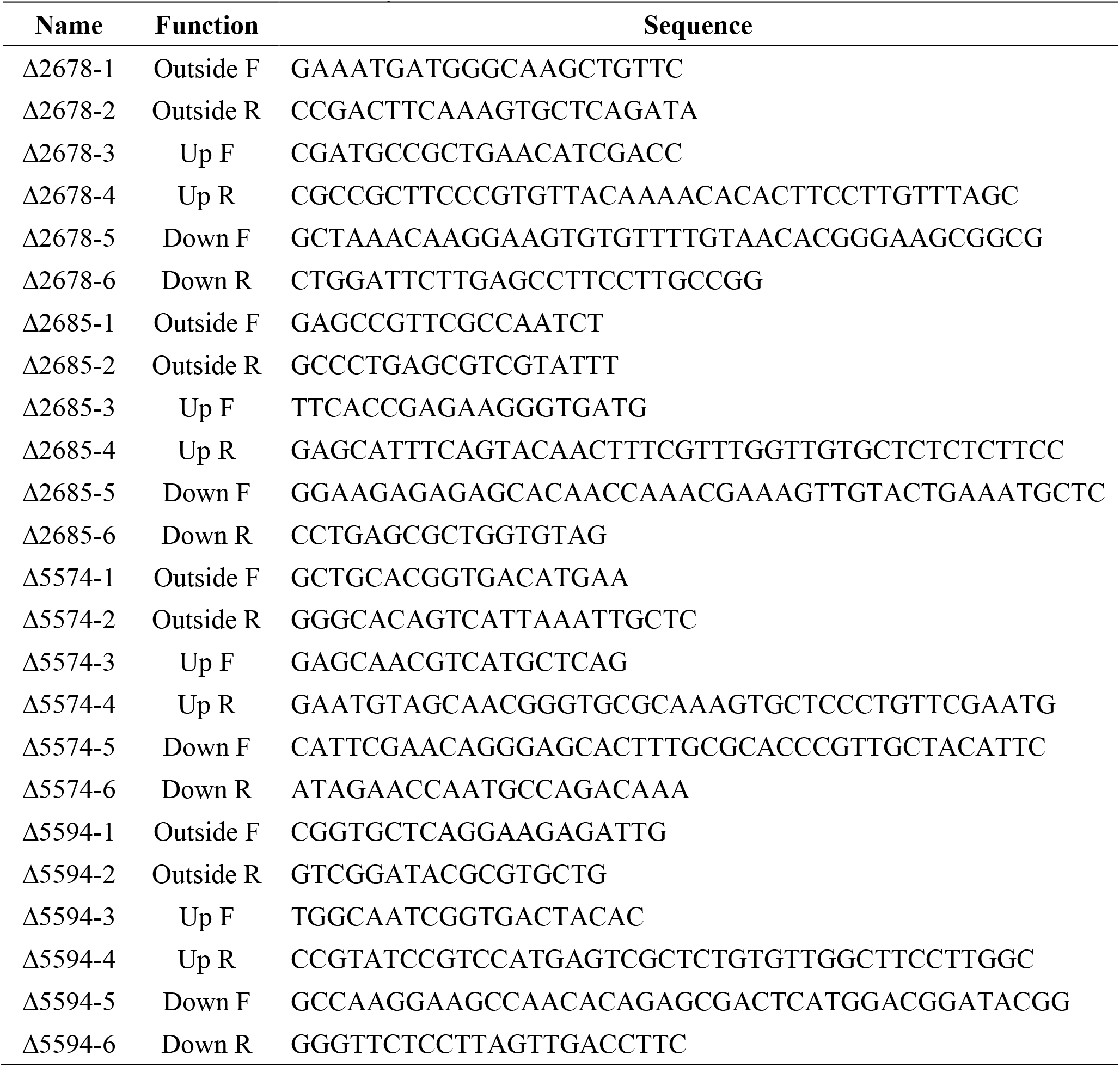
Primers used in this study.

**Table S3.** All proteins identified by LC-MS/MS.

| # | Locus | Identified Protein<sup>a</sup> | Spectrum Counts |  | Protein Probability |  |
| --- | --- | --- | --- | --- | --- | --- |
| | | | $\Delta rsmE$ | WT | $\Delta rsmE$ | WT |
| 1 | Pfl01_2678 | M10 family metallopeptidase | 193 | 0 | 100% | 0% |
| 2 | Pfl01_1917 | L-asparaginase, type II | 67 | 0 | 100% | 0% |
| 3 | Pfl01_0849 | Fumarate hydratase | 64 | 0 | 100% | 0% |
| 4 | Pfl01_2064 | Lysine-arginine-ornithine-binding periplasmic protein | 62 | 0 | 100% | 0% |
| 5 | Pfl01_4545 | 5-methyltetrahydropteroyltriglutamate-- homocysteine S-methyltransferase | 45 | 0 | 100% | 0% |
| 6 | Pfl01_3363 | M10 family metallopeptidase | 47 | 0 | 100% | 0% |
| 7 | Pfl01_2563 | Peptidase S45, penicillin amidase | 40 | 0 | 100% | 0% |
| 8 | Pfl01_2868 | Thiolase | 38 | 0 | 100% | 0% |
| 9 | Pfl01_0851 | Manganese and iron superoxide dismutase | 37 | 0 | 100% | 0% |
| 10 | Pfl01_2074 | Acetyl-CoA acetyltransferase | 36 | 0 | 100% | 0% |
| 11 | Pfl01_5072 | 50S ribosomal protein L16 | 34 | 0 | 100% | 0% |
| 12 | Pfl01_5265 | Phosphoglycerate kinase | 34 | 0 | 100% | 0% |
| 13 | Pfl01_0336 | Preprotein translocase subunit SecB | 33 | 0 | 100% | 0% |
| 14 | Pfl01_4062 | DegT/DnrJ/EryC1/StrS aminotransferase | 30 | 0 | 100% | 0% |
| 15 | Pfl01_1893 | Aspartate-semialdehyde dehydrogenase | 25 | 0 | 100% | 0% |
| 16 | Pfl01_0268 | Phosphoenolpyruvate carboxykinase | 22 | 0 | 100% | 0% |
| 17 | Pfl01_0333 | Phosphoglyceromutase | 22 | 0 | 100% | 0% |
| 18 | Pfl01_0618 | Acetyl-CoA carboxylase biotin carboxylase subunit | 21 | 0 | 100% | 0% |
| 19 | Pfl01_3879 | Multifunctional fatty acid oxidation complex subunit alpha | 18 | 0 | 100% | 0% |
| 20 | Pfl01_1248 | Extracellular ligand-binding receptor | 23 | 0 | 100% | 0% |
| 21 | Pfl01_4695 | Aldo/keto reductase | 22 | 0 | 100% | 0% |
| 22 | Pfl01_1625 | Heat shock protein 90 | 20 | 0 | 100% | 0% |
| 23 | Pfl01_1413 | Cysteine synthase | 17 | 0 | 100% | 0% |
| 24 | Pfl01_4361 | Keto-hydroxyglutarate-aldolase/keto-deoxy- phosphogluconate aldolase | 18 | 0 | 100% | 0% |
| 25 | Pfl01_4587 | GMP synthase | 18 | 0 | 100% | 0% |
| 26 | Pfl01_2925 | Alkyl hydroperoxide reductase | 20 | 0 | 100% | 0% |
| 27 | Pfl01_1022 | 50S ribosomal protein L19 | 17 | 0 | 100% | 0% |
| 28 | Pfl01_1311 | Iron-containing alcohol dehydrogenase | 16 | 0 | 100% | 0% |
| 29 | Pfl01_1852 | Twin-arginine translocation pathway signal protein | 17 | 0 | 100% | 0% |
| 30 | Pfl01_4190 | Hypothetical protein | 19 | 0 | 100% | 0% |
| 31 | Pfl01_4314 | Glycerophosphodiester phosphodiesterase | 16 | 0 | 100% | 0% |
| 32 | Pfl01_5574 | Type VI secretion system tip protein VgrG | 12 | 0 | 100% | 0% |
| 33 | Pfl01_0090 | Oligopeptidase A | 16 | 0 | 100% | 0% |
| 34 | Pfl01_2329 | Rhs element Vgr protein | 16 | 0 | 100% | 0% |
| 35 | Pfl01_3927 | Periplasmic solute binding protein | 15 | 0 | 100% | 0% |
| 36 | Pfl01_5511 | Methyltransferase | 15 | 0 | 100% | 0% |
| 37 | Pfl01_4376 | Glyceraldehyde-3-phosphate dehydrogenase | 15 | 0 | 100% | 0% |
| 38 | Pfl01_2866 | Short chain enoyl-CoA hydratase | 15 | 0 | 100% | 0% |
| 39 | Pfl01_5078 | 50S ribosomal protein L4 | 17 | 0 | 100% | 0% |
| 40 | Pfl01_1100 | Methionine aminopeptidase | 12 | 0 | 100% | 0% |
| 41 | Pfl01_4813 | Glucose-6-phosphate isomerase | 12 | 0 | 100% | 0% |
| 42 | Pfl01_2869 | Short-chain dehydrogenase/reductase SDR | 14 | 0 | 100% | 0% |
| 43 | Pfl01_5077 | 50S ribosomal protein L23 | 13 | 0 | 100% | 0% |
| 44 | Pfl01_4292 | Lysine-arginine-ornithine-binding periplasmic protein | 7 | 0 | 100% | 0% |
| 45 | Pfl01_3695 | Lon-A peptidase | 14 | 0 | 100% | 0% |
| 46 | Pfl01_0344 | L-glutamine synthetase | 12 | 0 | 100% | 0% |
| 47 | Pfl01_1925 | Inosine/uridine-preferring nucleoside hydrolase | 12 | 0 | 100% | 0% |
| 48 | Pfl01_3640 | Cyclophilin type peptidyl-prolyl cis-trans isomerase | 12 | 0 | 100% | 0% |
| 49 | Pfl01_5102 | N-acetyl-gamma-glutamyl-phosphate reductase | 11 | 0 | 100% | 0% |
| 50 | Pfl01_1849 | Amino acid adenylation protein | 12 | 0 | 100% | 0% |
| 51 | Pfl01_0612 | Phosphoribosylamine--glycine ligase | 12 | 0 | 100% | 0% |
| 52 | Pfl01_2212 | Amino acid adenylation protein | 4 | 0 | 100% | 0% |
| 53 | Pfl01_5731 | F0F1 ATP synthase subunit gamma | 10 | 0 | 100% | 0% |
| 54 | Pfl01_3939 | Alpha/beta hydrolase | 11 | 0 | 100% | 0% |
| 55 | Pfl01_2865 | Acyl-CoA dehydrogenase | 11 | 0 | 100% | 0% |
| 56 | Pfl01_1160 | Phage baseplate assembly protein V | 10 | 0 | 100% | 0% |
| 57 | Pfl01_4045 | Hexapptide repeat-containing transferase | 11 | 0 | 100% | 0% |
| 58 | Pfl01_1499 | Phenylalanine 4-monooxygenase | 10 | 0 | 100% | 0% |
| 59 | Pfl01_5049 | Single-stranded DNA-binding protein | 11 | 0 | 100% | 0% |
| 60 | Pfl01_4380 | Heme oxygenase | 10 | 0 | 100% | 0% |
| 61 | Pfl01_1572 | CheW protein | 6 | 0 | 100% | 0% |
| 62 | Pfl01_5621 | Phosphoribosylaminoimidazole carboxylase ATPase subunit | 9 | 0 | 100% | 0% |
| 63 | Pfl01_2044 | Rhs element Vgr protein | 7 | 0 | 100% | 0% |
| 64 | Pfl01_2192 | NADPH-dependent FMN reductase | 9 | 0 | 100% | 0% |
| 65 | Pfl01_5572 | Hypothetical protein | 10 | 0 | 100% | 0% |
| 66 | Pfl01_1764 | 2-methylisocitrate lyase | 9 | 0 | 100% | 0% |
| 67 | Pfl01_1768 | 2-methylcitrate dehydratase | 8 | 0 | 100% | 0% |
| 68 | Pfl01_5381 | Threonine dehydratase | 8 | 0 | 100% | 0% |
| 69 | Pfl01_5594 | Type VI secretion system contractile sheath large subunit | 9 | 0 | 100% | 0% |
| 70 | Pfl01_5043 | Hypothetical protein | 6 | 0 | 100% | 0% |
| 71 | Pfl01_5524 | Alanine racemase | 8 | 0 | 100% | 0% |
| 72 | Pfl01_0836 | Aspartyl/glutamyl-tRNA amidotransferase subunit A | 8 | 0 | 100% | 0% |
| 73 | Pfl01_2628 | Hypothetical protein | 5 | 0 | 100% | 0% |
| 74 | Pfl01_5498 | Diaminopimelate decarboxylase | 7 | 0 | 100% | 0% |
| 75 | Pfl01_0871 | Histidinol dehydrogenase | 8 | 0 | 100% | 0% |
| 76 | Pfl01_1498 | Pterin-4-alpha-carbinolamine dehydratase | 8 | 0 | 100% | 0% |
| 77 | Pfl01_5358 | Dihydroxy-acid dehydratase | 7 | 0 | 100% | 0% |
| 78 | Pfl01_0311 | Extracellular solute-binding protein | 6 | 0 | 100% | 0% |
| 79 | Pfl01_1104 | Ribosome recycling factor | 7 | 0 | 100% | 0% |
| 80 | Pfl01_4507 | Methionyl-tRNA synthetase | 6 | 0 | 100% | 0% |
| 81 | Pfl01_4281 | Succinylarginine dihydrolase | 5 | 0 | 100% | 0% |
| 82 | Pfl01_4362 | 6-phosphogluconolactonase | 6 | 0 | 100% | 0% |
| 83 | Pfl01_1185 | Oxidoreductase FAD/NAD(P)-binding | 7 | 0 | 100% | 0% |
| 84 | Pfl01_0366 | Histidine ammonia-lyase | 6 | 0 | 100% | 0% |
| 85 | Pfl01_1878 | Glutamyl-tRNA synthetase | 5 | 0 | 100% | 0% |
| 86 | Pfl01_4616 | Inositol monophosphatase | 4 | 0 | 100% | 0% |
| 87 | Pfl01_4093 | Fumarate hydratase | 6 | 0 | 100% | 0% |
| 88 | Pfl01_3466 | Dihydrolipoamide dehydrogenase | 5 | 0 | 100% | 0% |
| 89 | Pfl01_4046 | DegT/DnrJ/EryC1/StrS aminotransferase | 6 | 0 | 100% | 0% |
| 90 | Pfl01_1060 | Lysyl-tRNA synthetase | 5 | 0 | 100% | 0% |
| 91 | Pfl01_5246 | Hypothetical protein | 5 | 0 | 100% | 0% |
| 92 | Pfl01_4427 | Bifunctional ornithine acetyltransferase/N-acetylglutamate synthase | 6 | 0 | 100% | 0% |
| 93 | Pfl01_3107 | B12-dependent methionine synthase | 6 | 0 | 100% | 0% |
| 94 | Pfl01_3638 | Cysteinyl-tRNA synthetase | 5 | 0 | 100% | 0% |
| 95 | Pfl01_1297 | Glycoside hydrolase family protein | 3 | 0 | 100% | 0% |
| 96 | Pfl01_5098 | Tyrosyl-tRNA synthetase | 4 | 0 | 100% | 0% |
| 97 | Pfl01_3658 | Gamma-carboxygeranoyl-CoA hydratase | 5 | 0 | 100% | 0% |
| 98 | Pfl01_5489 | Argininosuccinate lyase | 5 | 0 | 100% | 0% |
| 99 | Pfl01_2267 | Amino acid adenylation protein | 4 | 0 | 100% | 0% |
| 100 | Pfl01_4989 | Aldehyde dehydrogenase | 5 | 0 | 100% | 0% |
| 101 | Pfl01_4274 | Aspartate kinase | 5 | 0 | 100% | 0% |
| 102 | Pfl01_4517 | Argininosuccinate synthase | 4 | 0 | 100% | 0% |
| 103 | Pfl01_2052 | 2,3-dimethylmalate lyase | 4 | 0 | 100% | 0% |
| 104 | Pfl01_4279 | Succinylglutamate desuccinylase | 5 | 0 | 100% | 0% |
| 105 | Pfl01_3318 | TonB-dependent siderophore receptor | 3 | 0 | 100% | 0% |
| 106 | Pfl01_1679 | C-terminal processing peptidase-1 | 4 | 0 | 100% | 0% |
| 107 | Pfl01_5618 | Aldose 1-epimerase | 4 | 0 | 100% | 0% |
| 108 | Pfl01_1066 | OmpA/MotB | 4 | 0 | 100% | 0% |
| 109 | Pfl01_5131 | Aminoglycoside phosphotransferase | 3 | 0 | 100% | 0% |
| 110 | Pfl01_3886 | Erythronate-4-phosphate dehydrogenase | 4 | 0 | 100% | 0% |
| 111 | Pfl01_5542 | Phosphomannomutase | 4 | 0 | 100% | 0% |
| 112 | Pfl01_4974 | Gamma-glutamyl phosphate reductase | 3 | 0 | 100% | 0% |
| 113 | Pfl01_2928 | Glutathione reductase | 3 | 0 | 100% | 0% |
| 114 | Pfl01_5053 | Catalase-like protein | 3 | 0 | 100% | 0% |
| 115 | Pfl01_0010 | Glycyl-tRNA synthetase subunit alpha | 3 | 0 | 100% | 0% |
| 116 | Pfl01_5330 | Thiazole synthase | 3 | 0 | 100% | 0% |
| 117 | Pfl01_2583 | TonB-dependent siderophore receptor | 3 | 0 | 100% | 0% |
| 118 | Pfl01_5169 | Hypothetical protein | 3 | 0 | 100% | 0% |
| 119 | Pfl01_5208 | Serine hydroxymethyltransferase | 0 | 0 | 8% | 0% |
| 120 | Pfl01_3257 | Amidase | 4 | 0 | 100% | 5% |
| 121 | Pfl01_1892 | 3-isopropylmalate dehydrogenase | 11 | 0 | 100% | 6% |
| 122 | Pfl01_4092 | Hypothetical protein | 9 | 0 | 100% | 6% |
| 123 | Pfl01_3465 | Branched-chain alpha-keto acid dehydrogenase subunit E2 | 9 | 0 | 100% | 6% |
| 124 | Pfl01_3657 | Propionyl-CoA carboxylase | 3 | 0 | 100% | 6% |
| 125 | Pfl01_4391 | Glycine cleavage system protein H | 19 | 0 | 100% | 7% |
| 126 | Pfl01_2076 | 3-oxoacid CoA-transferase | 18 | 0 | 100% | 7% |
| 127 | Pfl01_5552 | Endoribonuclease L-PSP | 14 | 0 | 100% | 7% |
| 128 | Pfl01_1282 | NADH:flavin oxidoreductase/NADH oxidase | 7 | 0 | 100% | 7% |
| 129 | Pfl01_5262 | Fructose-1,6-bisphosphate aldolase | 3 | 0 | 100% | 7% |
| 130 | Pfl01_5506 | Nitrogen regulatory protein P-II | 9 | 0 | 100% | 7% |
| 131 | Pfl01_0527 | ATP phosphoribosyltransferase | 8 | 0 | 100% | 7% |
| 132 | Pfl01_1158 | DNA circulation-like | 6 | 0 | 100% | 7% |
| 133 | Pfl01_1038 | Glucan biosynthesis protein D | 5 | 0 | 100% | 7% |
| 134 | Pfl01_5730 | F0F1 ATP synthase subunit beta | 93 | 0 | 100% | 8% |
| 135 | Pfl01_0767 | Carbamoyl phosphate synthase large subunit | 9 | 0 | 100% | 8% |
| 136 | Pfl01_1851 | Hypothetical protein | 10 | 0 | 100% | 8% |
| 137 | Pfl01_4109 | Hypothetical protein | 8 | 0 | 100% | 8% |
| 138 | Pfl01_0592 | Periplasmic binding protein | 3 | 0 | 100% | 8% |
| 139 | Pfl01_3791 | Filamentous hemagglutinin-like protein | 4 | 0 | 100% | 8% |
| 140 | Pfl01_1765 | Methylcitrate synthase | 4 | 0 | 100% | 9% |
| 141 | Pfl01_1847 | Amino acid adenylation protein | 78 | 0 | 100% | 10% |
| 142 | Pfl01_4693 | 30S ribosomal protein S9 | 13 | 0 | 100% | 10% |
| 143 | Pfl01_1516 | Motility/biofilm formation protein FlhH | 5 | 0 | 100% | 10% |
| 144 | Pfl01_4598 | Pyrrolo-quinoline quinone | 6 | 0 | 100% | 10% |
| 145 | Pfl01_4786 | Ketol-acid reductoisomerase | 26 | 0 | 100% | 11% |
| 146 | Pfl01_0911 | Fumarylacetoacetate hydrolase | 28 | 0 | 100% | 11% |
| 147 | Pfl01_3847 | Transaldolase B | 21 | 0 | 100% | 11% |
| 148 | Pfl01_3656 | Isovaleryl-CoA dehydrogenase | 11 | 0 | 100% | 11% |
| 149 | Pfl01_1027 | Homoserine dehydrogenase | 5 | 0 | 100% | 11% |
| 150 | Pfl01_2631 | Rhs element Vgr protein | 21 | 0 | 100% | 13% |
| 151 | Pfl01_2685 | Triacylglycerol lipase | 6 | 0 | 100% | 13% |
| 152 | Pfl01_1110 | Outer membrane chaperone Skp | 6 | 0 | 100% | 13% |
| 153 | Pfl01_0360 | Urocanate hydratase | 4 | 0 | 100% | 13% |
| 154 | Pfl01_3602 | Isocitrate lyase | 20 | 0 | 100% | 14% |
| 155 | Pfl01_1612 | Succinate dehydrogenase flavoprotein subunit | 11 | 0 | 100% | 14% |
| 156 | Pfl01_5386 | FAD linked oxidase-like protein | 6 | 0 | 100% | 14% |
| 157 | Pfl01_2330 | RHS protein | 4 | 0 | 100% | 14% |
| 158 | Pfl01_3911 | Peptidase M24 | 21 | 0 | 100% | 15% |
| 159 | Pfl01_1148 | Phage tail tape measure protein TP901, core region | 5 | 0 | 100% | 15% |
| 160 | Pfl01_1154 | Bacteriophage Mu tail sheath | 53 | 0 | 100% | 16% |
| 161 | Pfl01_2045 | Hcp family type VI secretion system effector | 27 | 0 | 100% | 16% |
| 162 | Pfl01_5113 | Indole-3-glycerol-phosphate synthase | 9 | 0 | 100% | 16% |
| 163 | Pfl01_2882 | Enoyl-CoA hydratase | 4 | 0 | 100% | 16% |
| 164 | Pfl01_5002 | Hypothetical protein | 3 | 0 | 100% | 16% |
| 165 | Pfl01_4852 | Isoleucyl-tRNA synthetase | 19 | 0 | 100% | 17% |
| 166 | Pfl01_1609 | Type II citrate synthase | 12 | 0 | 100% | 17% |
| 167 | Pfl01_1497 | Aromatic amino acid aminotransferase | 57 | 0 | 100% | 18% |
| 168 | Pfl01_4158 | ACP S-malonyltransferase | 11 | 0 | 100% | 18% |
| 169 | Pfl01_4071 | NAD-dependent epimerase/dehydratase | 8 | 0 | 100% | 18% |
| 170 | Pfl01_0352 | Fructose-1,6-bisphosphatase | 10 | 0 | 100% | 19% |
| 171 | Pfl01_0133 | Von Willebrand factor A | 4 | 0 | 100% | 19% |
| 172 | Pfl01_2917 | 4-hydroxyphenylpyruvate dioxygenase | 10 | 0 | 100% | 20% |
| 173 | Pfl01_4860 | 50S ribosomal protein L21 | 19 | 0 | 100% | 22% |
| 174 | Pfl01_2200 | Aminopeptidase | 10 | 0 | 100% | 22% |
| 175 | Pfl01_4275 | Alanyl-tRNA synthetase | 9 | 0 | 100% | 22% |
| 176 | Pfl01_0986 | Valyl-tRNA synthetase | 6 | 0 | 100% | 22% |
| 177 | Pfl01_1919 | Periplasmic binding protein/LacI transcriptional regulator | 9 | 0 | 100% | 25% |
| 178 | Pfl01_5726 | Glucosamine--fructose-6-phosphate aminotransferase | 3 | 0 | 100% | 25% |
| 179 | Pfl01_2213 | Amino acid adenylation protein | 2 | 0 | 100% | 26% |
| 180 | Pfl01_0783 | Polynucleotide phosphorylase | 53 | 0 | 100% | 27% |
| 181 | Pfl01_4074 | Bifunctional cyclohexadienyl dehydrogenase/ 3-phosphoshikimate 1-carboxyvinyltransferase | 15 | 0 | 100% | 30% |
| 182 | Pfl01_4699 | Tryptophanyl-tRNA synthetase | 6 | 0 | 100% | 30% |
| 183 | Pfl01_0835 | Aspartyl/glutamyl-tRNA amidotransferase subunit B | 11 | 0 | 100% | 32% |
| 184 | Pfl01_4370 | Extracellular solute-binding protein | 3 | 0 | 100% | 32% |
| 185 | Pfl01_4394 | Glycine cleavage system T protein | 28 | 0 | 100% | 33% |
| 186 | Pfl01_5430 | Extracellular solute-binding protein | 16 | 0 | 100% | 33% |
| 187 | Pfl01_1141 | Baseplate J-like protein | 18 | 0 | 100% | 35% |
| 188 | Pfl01_5538 | Extracellular solute-binding protein | 6 | 0 | 100% | 35% |
| 189 | Pfl01_1145 | Phage tail sheath protein FI-like | 82 | 0 | 100% | 37% |
| 190 | Pfl01_1616 | Dihydrolipoamide dehydrogenase | 105 | 0 | 100% | 38% |
| 191 | Pfl01_0399 | Arginyl-tRNA synthetase | 12 | 0 | 100% | 38% |
| 192 | Pfl01_4957 | Leucyl-tRNA synthetase | 11 | 0 | 100% | 38% |
| 193 | Pfl01_3862 | Soluble pyridine nucleotide transhydrogenase | 5 | 0 | 100% | 38% |
| 194 | Pfl01_0462 | Dihydrolipoamide acetyltransferase | 3 | 0 | 100% | 38% |
| 195 | Pfl01_4295 | Ribonucleotide-diphosphate reductase subunit beta | 3 | 0 | 100% | 38% |
| 196 | Pfl01_4386 | Arginine deiminase | 63 | 0 | 100% | 41% |
| 197 | Pfl01_0052 | DSBA oxidoreductase | 9 | 0 | 100% | 41% |
| 198 | Pfl01_0295 | Outer membrane porin | 5 | 0 | 100% | 41% |
| 199 | Pfl01_4414 | Ferritin and Dps | 27 | 0 | 100% | 42% |
| 200 | Pfl01_4532 | Glycerol kinase | 59 | 1 | 100% | 44% |
| 201 | Pfl01_5074 | 50S ribosomal protein L22 | 8 | 1 | 100% | 44% |
| 202 | Pfl01_4519 | Dihydroorotase | 7 | 1 | 100% | 44% |
| 203 | Pfl01_5280 | S-adenosyl-L-homocysteine hydrolase | 49 | 1 | 100% | 47% |
| 204 | Pfl01_3639 | Glutaminyl-tRNA synthetase | 23 | 0 | 100% | 47% |
| 205 | Pfl01_1166 | Hypothetical protein | 15 | 1 | 100% | 47% |
| 206 | Pfl01_4588 | Inosine 5'-monophosphate dehydrogenase | 6 | 1 | 100% | 47% |
| 207 | Pfl01_4401 | OmpA/MotB | 82 | 0 | 100% | 49% |
| 208 | Pfl01_1164 | Hypothetical protein | 35 | 0 | 100% | 51% |
| 209 | Pfl01_4411 | Aspartyl-tRNA synthetase | 23 | 1 | 100% | 52% |
| 210 | Pfl01_0188 | 4-aminobutyrate aminotransferase | 42 | 0 | 100% | 53% |
| 211 | Pfl01_3699 | Bifunctional 5,10-methylene-tetrahydrofolate dehydrogenase/5,10-methylene-tetrahydrofolate cyclohydrolase | 4 | 1 | 100% | 56% |
| 212 | Pfl01_0596 | TonB-dependent copper receptor | 19 | 1 | 100% | 57% |
| 213 | Pfl01_5457 | Thioredoxin | 35 | 1 | 100% | 58% |
| 214 | Pfl01_5058 | 30S ribosomal protein S13 | 15 | 1 | 100% | 58% |
| 215 | Pfl01_5183 | Malate synthase G | 8 | 0 | 100% | 58% |
| 216 | Pfl01_4488 | Hypothetical protein | 83 | 1 | 100% | 59% |
| 217 | Pfl01_1591 | Thiosulfate-binding protein | 4 | 0 | 100% | 59% |
| 218 | Pfl01_0613 | Bifunctional phosphoribosylaminoimidazolecarboxamide formyltransferase/IMP cyclohydrolase | 21 | 0 | 100% | 60% |
| 219 | Pfl01_3395 | Bifunctional aconitate hydratase 2/2-methylisocitrate dehydratase | 28 | 0 | 100% | 61% |
| 220 | Pfl01_4117 | Electron transfer flavoprotein subunit beta | 44 | 0 | 100% | 62% |
| 221 | Pfl01_1371 | Dihydrodipicolinate synthase | 12 | 1 | 100% | 62% |
| 222 | Pfl01_5267 | Transketolase | 16 | 0 | 100% | 63% |
| 223 | Pfl01_1846 | Peptide synthase | 36 | 0 | 100% | 64% |
| 224 | Pfl01_5534 | Aldehyde dehydrogenase | 18 | 1 | 100% | 64% |
| 225 | Pfl01_2670 | Phosphoglucomutase | 15 | 0 | 100% | 64% |
| 226 | Pfl01_1506 | Flagellar hook-associated protein FlgK | 79 | 0 | 100% | 65% |
| 227 | Pfl01_1014 | Phosphoribosylglycinamide formyltransferase 2 | 9 | 0 | 100% | 66% |
| 228 | Pfl01_5284 | Hypothetical protein | 8 | 0 | 100% | 66% |
| 229 | Pfl01_0528 | Adenylosuccinate synthetase | 33 | 0 | 100% | 67% |
| 230 | Pfl01_2270 | TauD/TfdA family dioxygenase | 11 | 1 | 100% | 67% |
| 231 | Pfl01_1070 | Adenylate kinase | 39 | 0 | 100% | 69% |
| 232 | Pfl01_5405 | Extracellular solute-binding protein | 59 | 1 | 100% | 71% |
| 233 | Pfl01_0813 | Extracellular solute-binding protein | 32 | 0 | 100% | 71% |
| 234 | Pfl01_1845 | Peptide synthase | 96 | 0 | 100% | 72% |
| 235 | Pfl01_1770 | Phosphoenolpyruvate synthase | 63 | 0 | 100% | 74% |
| 236 | Pfl01_4285 | Bifunctional N-succinyldiaminopimelate-aminotransferase/acetylornithine transaminase | 11 | 1 | 100% | 74% |
| 237 | Pfl01_3865 | Glyceraldehyde-3-phosphate dehydrogenase | 3 | 0 | 99% | 74% |
| 238 | Pfl01_5054 | 50S ribosomal protein L17 | 35 | 0 | 100% | 75% |
| 239 | Pfl01_0492 | Thiamine biosynthesis protein ThiC | 16 | 0 | 100% | 75% |
| 240 | Pfl01_0463 | Pyruvate dehydrogenase subunit E1 | 79 | 0 | 100% | 77% |
| 241 | Pfl01_0557 | Short-chain dehydrogenase/reductase SDR | 14 | 1 | 100% | 77% |
| 242 | Pfl01_4387 | Ornithine carbamoyltransferase | 67 | 0 | 100% | 78% |
| 243 | Pfl01_5073 | 30S ribosomal protein S3 | 6 | 1 | 100% | 78% |
| 244 | Pfl01_1934 | 50S ribosomal protein L20 | 30 | 1 | 100% | 80% |
| 245 | Pfl01_3940 | Peptide synthase | 54 | 1 | 100% | 81% |
| 246 | Pfl01_3206 | Glutamate dehydrogenase (NAD) | 9 | 0 | 100% | 82% |
| 247 | Pfl01_4594 | 2-isopropylmalate synthase | 10 | 1 | 100% | 82% |
| 248 | Pfl01_3698 | Trigger factor | 9 | 0 | 100% | 82% |
| 249 | Pfl01_0546 | Azurin | 44 | 1 | 100% | 83% |
| 250 | Pfl01_3594 | Isocitrate dehydrogenase (NADP+) | 24 | 0 | 100% | 83% |
| 251 | Pfl01_5086 | DNA-directed RNA polymerase subunit beta | 55 | 0 | 100% | 84% |
| 252 | Pfl01_5639 | Pyruvate carboxylase subunit B | 34 | 1 | 100% | 84% |
| 253 | Pfl01_3593 | Isocitrate dehydrogenase (NADP) | 81 | 1 | 100% | 85% |
| 254 | Pfl01_5075 | 30S ribosomal protein S19 | 18 | 1 | 100% | 85% |
| 255 | Pfl01_4419 | Outer membrane porin | 19 | 2 | 100% | 86% |
| 256 | Pfl01_5069 | 50S ribosomal protein L14 | 29 | 1 | 100% | 87% |
| 257 | Pfl01_1019 | 30S ribosomal protein S16 | 9 | 2 | 100% | 87% |
| 258 | Pfl01_1143 | Phage tail Collar | 74 | 1 | 100% | 89% |
| 259 | Pfl01_0244 | Cystine transporter subunit | 44 | 0 | 100% | 89% |
| 260 | Pfl01_3999 | Elongation factor P | 21 | 1 | 100% | 89% |
| 261 | Pfl01_4395 | Cold-shock DNA-binding protein | 20 | 2 | 100% | 89% |
| 262 | Pfl01_5387 | D-3-phosphoglycerate dehydrogenase | 5 | 1 | 100% | 89% |
| 263 | Pfl01_1777 | OmpF | 54 | 1 | 100% | 90% |
| 264 | Pfl01_4448 | Pyruvate kinase | 27 | 1 | 100% | 90% |
| 265 | Pfl01_5056 | 30S ribosomal protein S4 | 17 | 1 | 100% | 90% |
| 266 | Pfl01_5090 | 50S ribosomal protein L11 | 5 | 2 | 100% | 90% |
| 267 | Pfl01_2177 | Enoyl-ACP reductase | 40 | 1 | 100% | 91% |
| 268 | Pfl01_5055 | DNA-directed RNA polymerase subunit alpha | 26 | 1 | 100% | 91% |
| 269 | Pfl01_4535 | Extracellular solute-binding protein | 73 | 1 | 100% | 92% |
| 270 | Pfl01_5085 | DNA-directed RNA polymerase subunit beta' | 63 | 2 | 100% | 92% |
| 271 | Pfl01_1121 | Phosphopyruvate hydratase | 73 | 1 | 100% | 93% |
| 272 | Pfl01_5067 | 50S ribosomal protein L5 | 36 | 1 | 100% | 94% |
| 273 | Pfl01_4521 | Alkyl hydroperoxide reductase | 72 | 1 | 100% | 95% |
| 274 | Pfl01_5063 | 50S ribosomal protein L18 | 30 | 2 | 100% | 96% |
| 275 | Pfl01_1929 | Cold-shock DNA-binding protein family protein | 17 | 2 | 100% | 96% |
| 276 | Pfl01_5064 | 50S ribosomal protein L6 | 11 | 2 | 100% | 96% |
| 277 | Pfl01_0782 | 30S ribosomal protein S15 | 5 | 1 | 100% | 96% |
| 278 | Pfl01_4392 | Glycine dehydrogenase | 80 | 1 | 100% | 97% |
| 279 | Pfl01_1614 | 2-oxoglutarate dehydrogenase E1 | 25 | 2 | 100% | 97% |
| 280 | Pfl01_0814 | Outer membrane porin | 11 | 2 | 100% | 97% |
| 281 | Pfl01_0534 | 30S ribosomal protein S6 | 9 | 3 | 100% | 97% |
| 282 | Pfl01_5084 | 30S ribosomal protein S12 | 10 | 2 | 100% | 97% |
| 283 | Pfl01_0403 | Malic enzyme | 84 | 3 | 100% | 98% |
| 284 | Pfl01_5083 | 30S ribosomal protein S7 | 60 | 2 | 100% | 98% |
| 285 | Pfl01_5088 | 50S ribosomal protein L10 | 45 | 2 | 100% | 98% |
| 286 | Pfl01_5089 | 50S ribosomal protein L1 | 14 | 3 | 100% | 98% |
| 287 | Pfl01_1168 | Tail fiber protein | 11 | 1 | 100% | 98% |
| 288 | Pfl01_4072 | 30S ribosomal protein S1 | 5 | 2 | 100% | 98% |
| 289 | Pfl01_5079 | 50S ribosomal protein L3 | 45 | 2 | 100% | 99% |
| 290 | Pfl01_1618 | Succinyl-CoA synthetase subunit alpha | 40 | 4 | 100% | 99% |
| 291 | Pfl01_5625 | Aspartate ammonia-lyase | 37 | 2 | 100% | 99% |
| 292 | Pfl01_0223 | NLPA lipoprotein | 35 | 2 | 100% | 99% |
| 293 | Pfl01_4379 | TonB-dependent hemoglobin/transferrin/lactoferrin receptor | 27 | 2 | 100% | 99% |
| 294 | Pfl01_1162 | Baseplate J-like protein | 9 | 2 | 100% | 99% |
| 295 | Pfl01_1101 | 30S ribosomal protein S2 | 7 | 3 | 100% | 99% |
| 296 | Pfl01_5417 | Outer membrane porin | 7 | 2 | 100% | 99% |
| 297 | Pfl01_5081 | Elongation factor Tu | 228 | 15 | 100% | 100% |
| 298 | Pfl01_5082 | Elongation factor G | 164 | 6 | 100% | 100% |
| 299 | Pfl01_4501 | Chaperonin GroEL | 147 | 1 | 100% | 100% |
| 300 | Pfl01_5732 | F0F1 ATP synthase subunit alpha | 124 | 3 | 100% | 100% |
| 301 | Pfl01_1617 | Succinyl-CoA synthetase subunit beta | 100 | 5 | 100% | 100% |
| 302 | Pfl01_4877 | Serine hydroxymethyltransferase | 77 | 2 | 100% | 100% |
| 303 | Pfl01_1527 | Flagellin | 64 | 6 | 100% | 100% |
| 304 | Pfl01_0763 | Molecular chaperone DnaK | 58 | 2 | 100% | 100% |
| 305 | Pfl01_1848 | TonB-dependent siderophore receptor | 46 | 3 | 100% | 100% |
| 306 | Pfl01_4605 | Nucleoside diphosphate kinase | 30 | 5 | 100% | 100% |
| 307 | Pfl01_1102 | Elongation factor Ts | 22 | 2 | 100% | 100% |
| 308 | Pfl01_1529 | Flagellar hook-associated 2-like | 12 | 3 | 100% | 100% |
| 309 | Pfl01_5029 | OmpW | 7 | 7 | 100% | 100% |
| 310 | Pfl01_4011 | Outer membrane lipoprotein OprI | 8 | 2 | 100% | 100% |

<sup>a</sup> Annotated function information is from the *Pseudomonas* database (52).

**Table S4.**
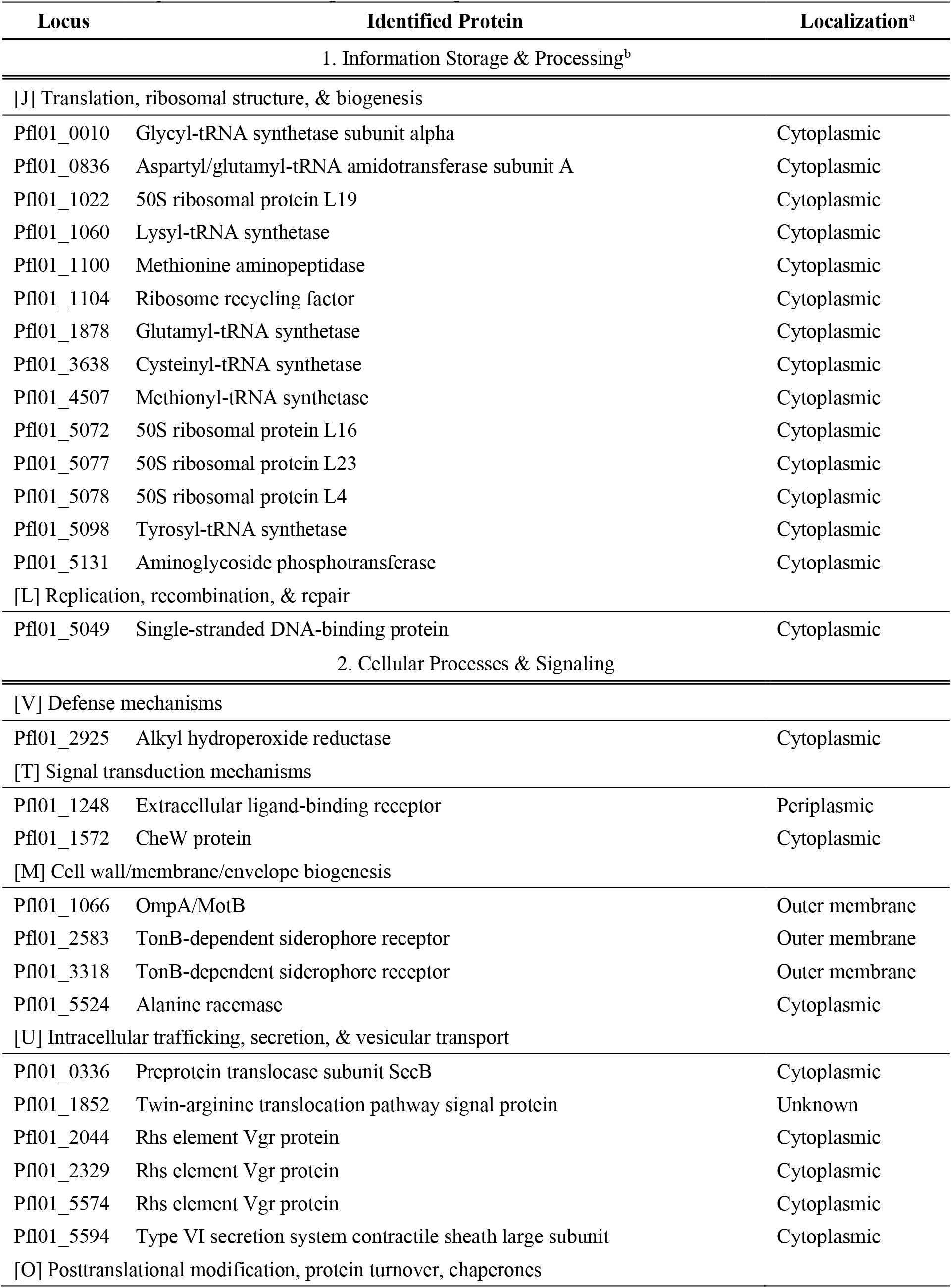
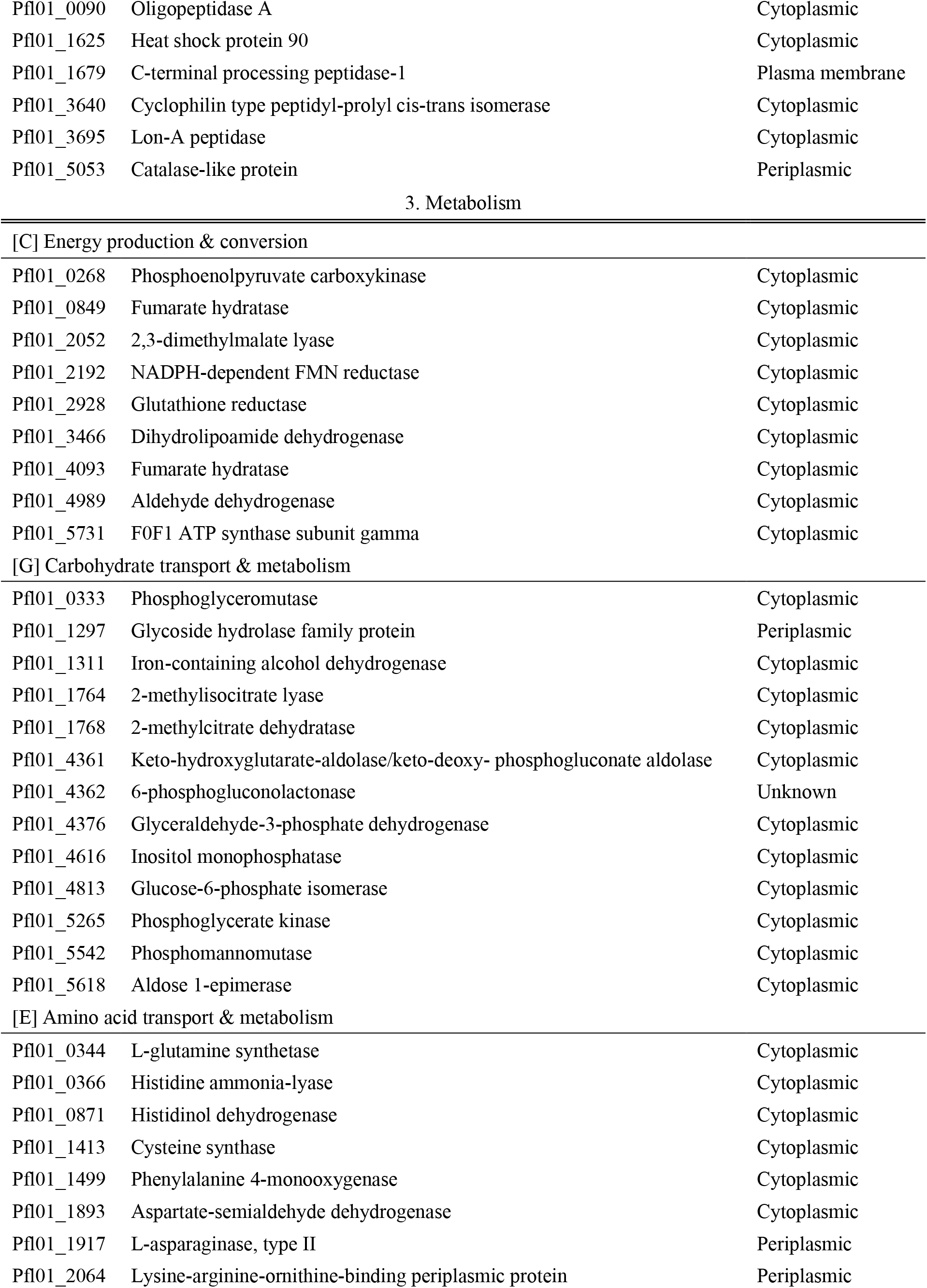
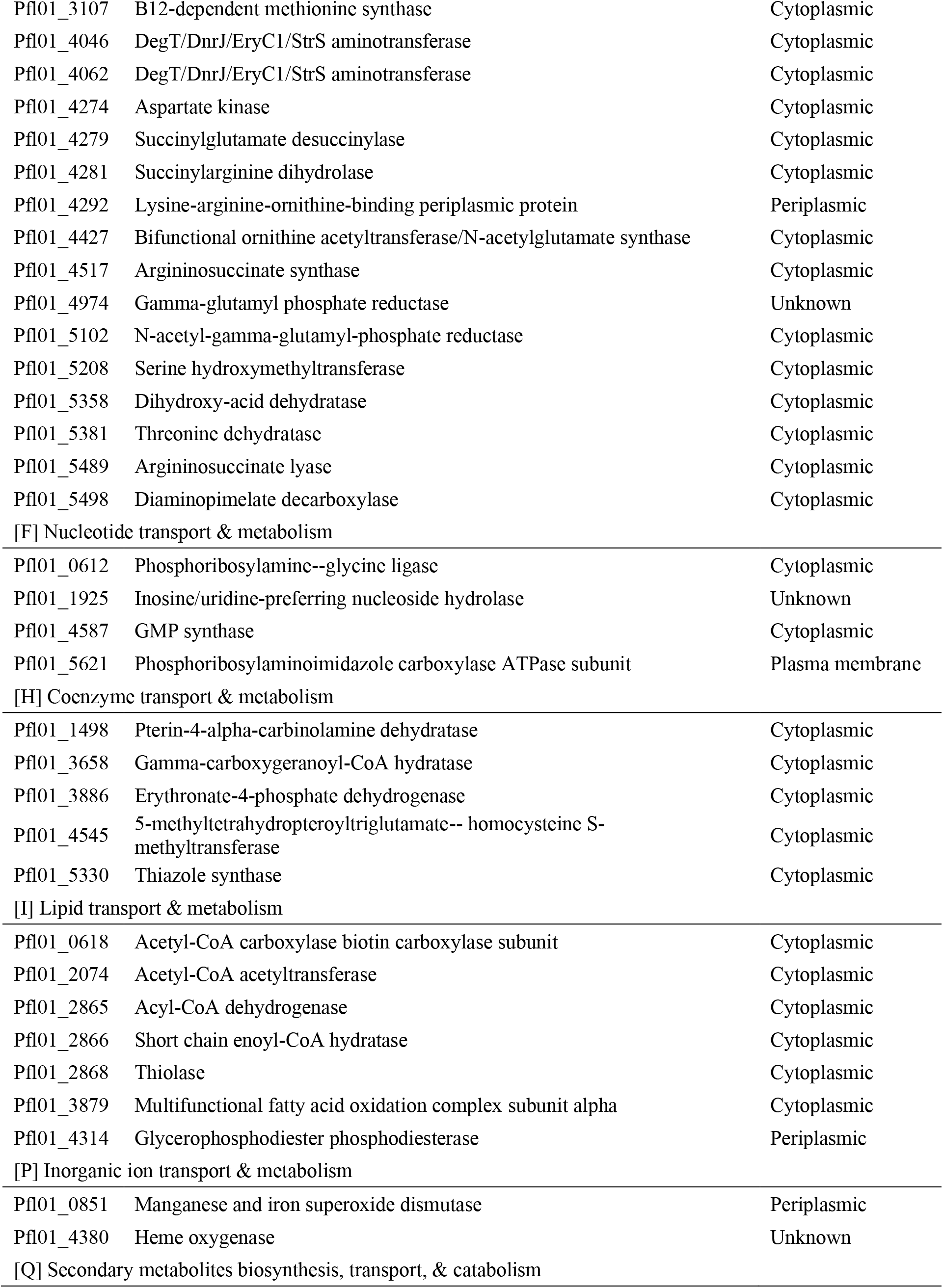
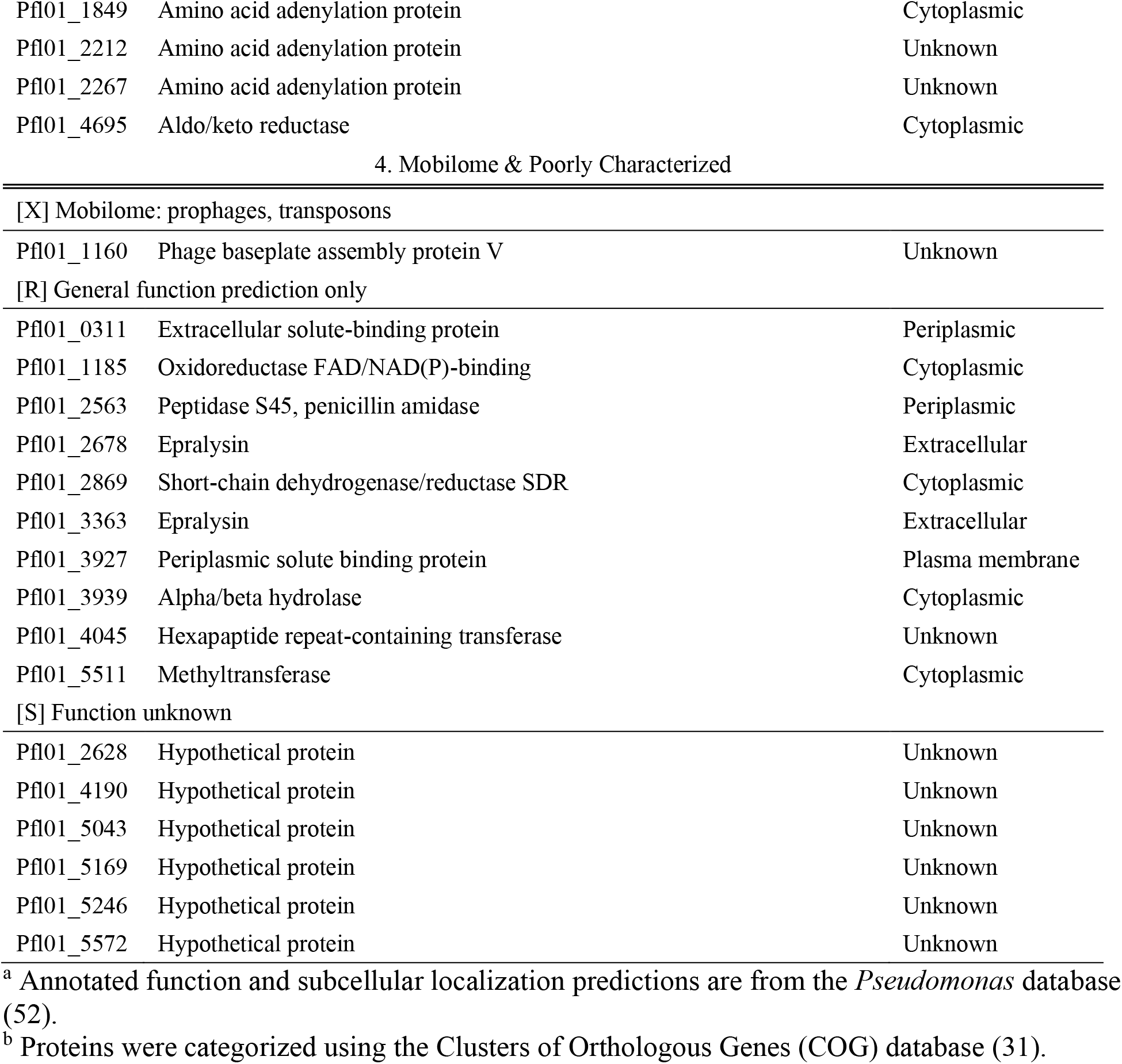
Categorization of 119 proteins unique to Δ*rsmE*.

**Figure S1.**
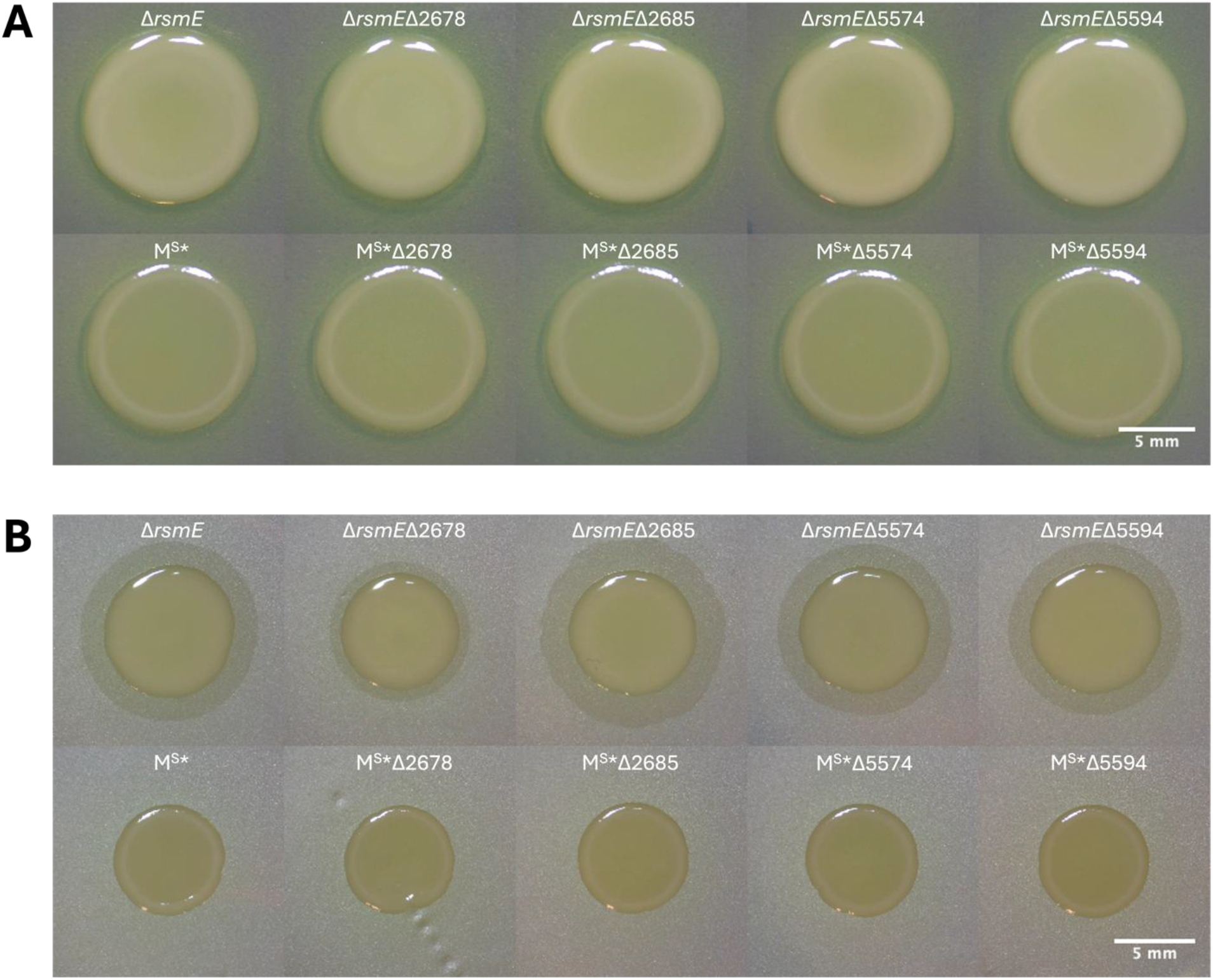
Knockouts of T6SS or extracellular enzyme genes do not affect extracellular polysaccharide or surfactant production. (A) Morphology of each mutant was imaged after one day of growth. All the mutants are similar in appearance to their parent strain, with the Δ*rsmE* appearing mucoid and M^S^* nonmucoid, so deleting the extracellular enzymes and T6SS genes does not alter extracellular polysaccharide production. (B) Surfactant screen of mutants spotted on the dull side of a polycarbonate membrane overlain on PAF was imaged after one day of growth. All the mutants in Δ*rsmE* background produce a surfactant ring like their parent isolate, and all the mutants in the M^S^* background do not produce a surfactant ring. Therefore, deleting the extracellular enzymes and T6SS genes does not affect biosurfactant production. Scale bars represent 5 mm.

**Figure S2.**
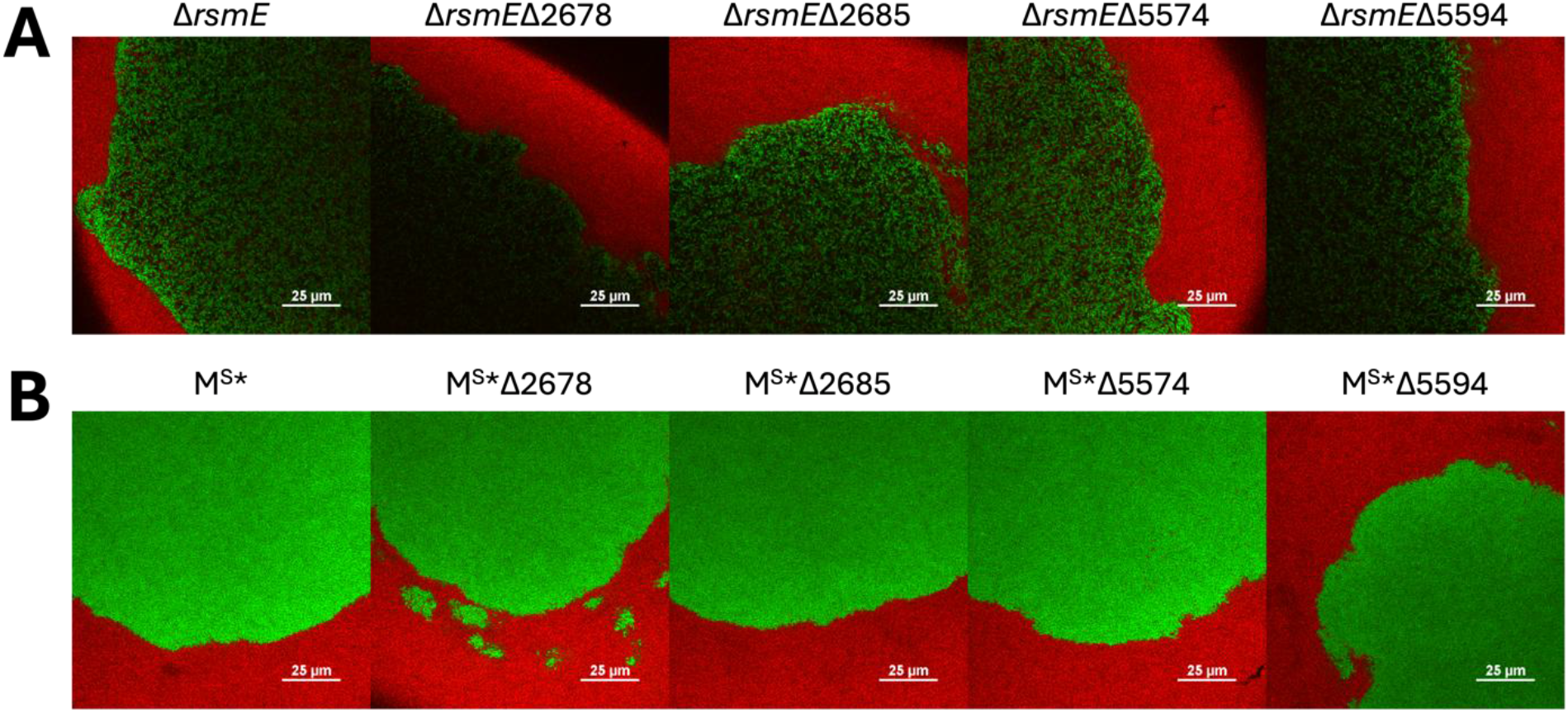
The T6SS is not a main driver of patch formation. GFP-labeled mutants were mixed with dsRedExpress-labeled WT at a 10^-5^ relative frequency and incubated for three days. Confocal images of the emergent patches created by the gene knockouts in (A) Δ*rsmE* and (B) M^S^* were obtained using the 100x TU PLAN APO objective. Deletion of the extracellular enzymes and the T6SS do not show changes in the size or general structure of the mutant patches. Scale bars represent 25 µm.

**Figure S3.**
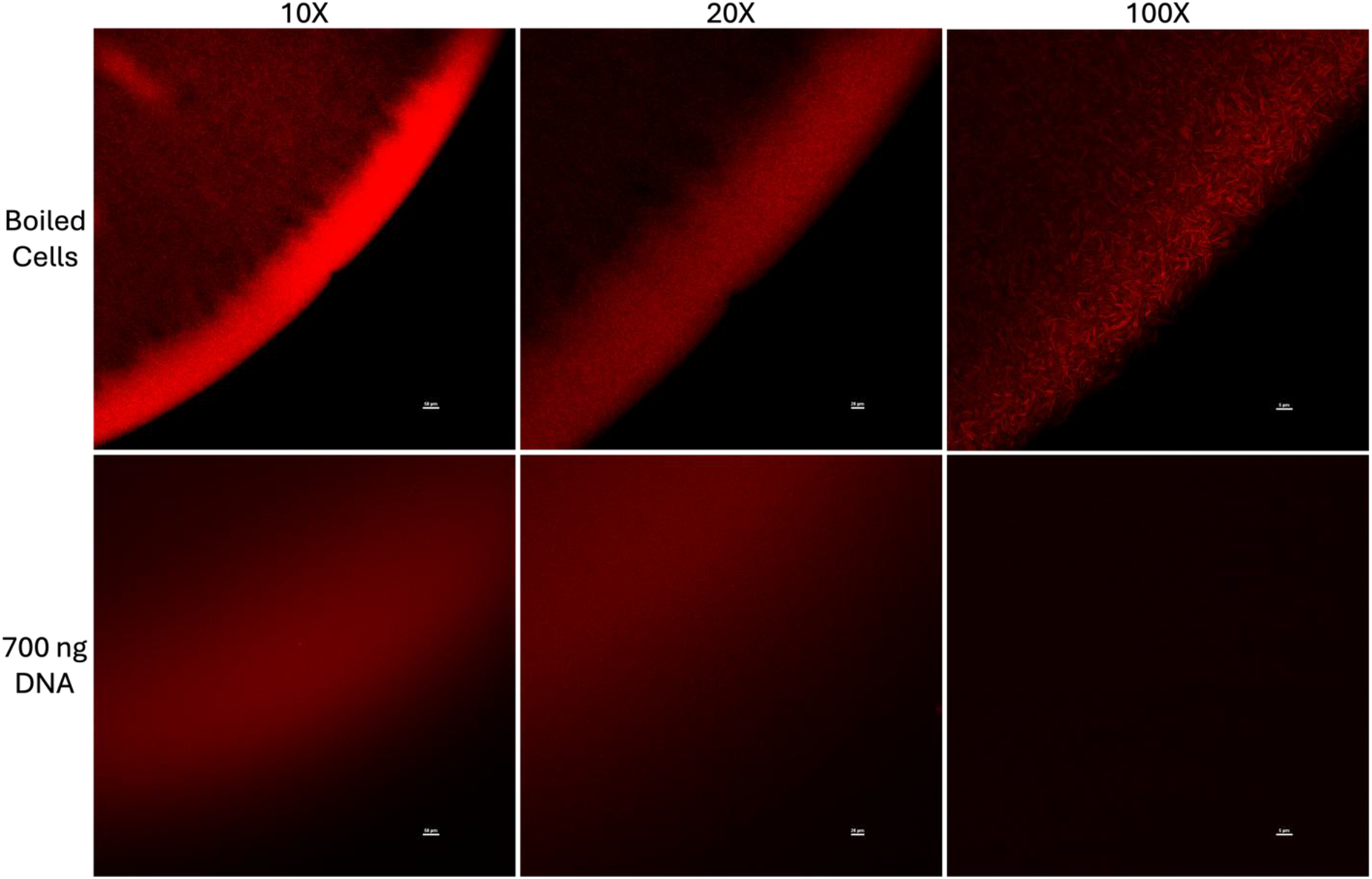
Propidium iodide (PI) signal from eDNA is only detectable using confocal microscopy at low magnifications. To image dead cells (top), WT cells in LB broth were boiled for 10 minutes, mixed with PI, and spotted on PAF. For eDNA (bottom), 700 ng of DNA was mixed with PI and spotted onto PAF. Images were captured via confocal microscopy after one day of incubation at room temperature. Confocal images were captured using the 10x PLAN APO, 20x PLAN APO, and 100x TU PLAN APO objectives. While PI signal from both boiled cells and eDNA are seen at 10x and 20x, PI signal from eDNA at 100x is no longer detectable. Scale bars represent 50 µm (10x), 20 µm (20x), and 5 µm (100x).

